# Repeatable phenotypic but not genetic response to selection on body size in the black soldier fly

**DOI:** 10.1101/2025.02.25.640052

**Authors:** Tomas N. Generalovic, Wenjun Zhou, Leia Chengcheng Zhao, Sam Leonard, Ian A. Warren, Miha Pipan, Chris D. Jiggins

## Abstract

Polygenic traits are expected to show high genetic redundancy and therefore low repeatability in the genomic response to selection. We tested this prediction by selecting for large body size in the black soldier fly (*Hermetia illucens*). Over three replicate experiments selected for large body size, we found a strong and repeatable phenotypic response, with a mean 15% increase in body size. Selected lines also increased in larval growth rate (+19%) and average protein content (+14%), suggesting that selection on large body size does not result in strong trade-offs. In contrast to the predictability of the phenotypic response across replicates, whole genome sequencing identified a highly polygenic and non-repeatable genomic response. We identified 120, 301 and 157 outlier genomic regions in the three replicates, but high redundancy with only four shared regions. Among 12 candidate genes found in these regions, the insulin-like receptor gene (*HiInR*) was confirmed as regulating larval growth using a CRISPR knockout experiment. In summary, polygenic quantitative traits show high genetic redundancy, even where the phenotypic response to selection is highly repeatable.

## Introduction

Studies of adaptive change typically either take a molecular genetic approach that focuses on allelic change at specific loci and often identifies a few alleles of large effect^1^, or alternatively use a quantitative genetics approach that focuses on the phenotype, and assumes a highly polygenic genetic control^2^. Exemplifying the former are traits such as butterfly wing patterns, controlled by just a handful of loci with high repeatability across divergent taxa^3,4^. At the other extreme, quantitative traits such as human height are influenced by many genes, with many thousands of SNPs even outside core pathways influencing the phenotype, sometimes termed an ‘omnigenic’ genetic architecture^5^. Key characteristics of such polygenic traits are high genetic redundancy and low repeatability^1^. In other words, many different loci can have similar phenotypic effects, and repeated selection on the same trait, even from a similar starting point can involve different loci in each case^6,7^. Barghi et al. have introduced the concept of ‘adaptive architecture’, as a contrast to ‘genetic architecture’, in which the identity of specific loci may vary between replicates, but there is a consistent architecture of the trait. Such polygenic traits are less tractable and have not been as well studied by evolutionary biologists as large-effect loci controlling oligogenic traits.

One well studied and highly polygenic trait is body size. In *Drosophila melanogaster*, body size has a strong influence on fitness^8^ and varies adaptively in association with climate. There are also complex relationships between body size and other traits such as longevity, fecundity and development time^9–13^. An increase in size can involve numerous physiological and morphological changes that influence both cell size and number^14–16^. Here, we focus on body size in the black soldier fly, *Hermetia illucens* L. (Stratiomyidae), which offers a tractable system to understand adaptive genetic architecture.

The black soldier fly forms the basis for a major insect livestock industry, in which nutrients are recovered from food waste and returned to the food chain^17^. *H. illucens* dominates this ‘Insects as Food and Feed’ industry due to its generalist diet and highly voracious feeding behaviour^18,19^. Growth rate and body size are therefore potentially important traits for this nascent industry. However, despite its economic importance, there has been little work on genetic improvement of *H. illucens*. Two previous studies have shown significant phenotypic responses to selection on body size over sixteen and three generations, respectively^20,21^. Here, we artificially selected for large pupal body size in a replicated experiment and used low-coverage whole genome sequencing to study the genetic response to this selection (Figure 1). We ask: (1) How repeatable is the phenotypic response to selection? (2) Are there phenotypic trade-offs of body size with other life history traits? (3) How repeatable is the genetic response to selection? (4) Can experimental evolution be used to identify specific genes controlling a highly polygenic trait?

**Figure 1.**
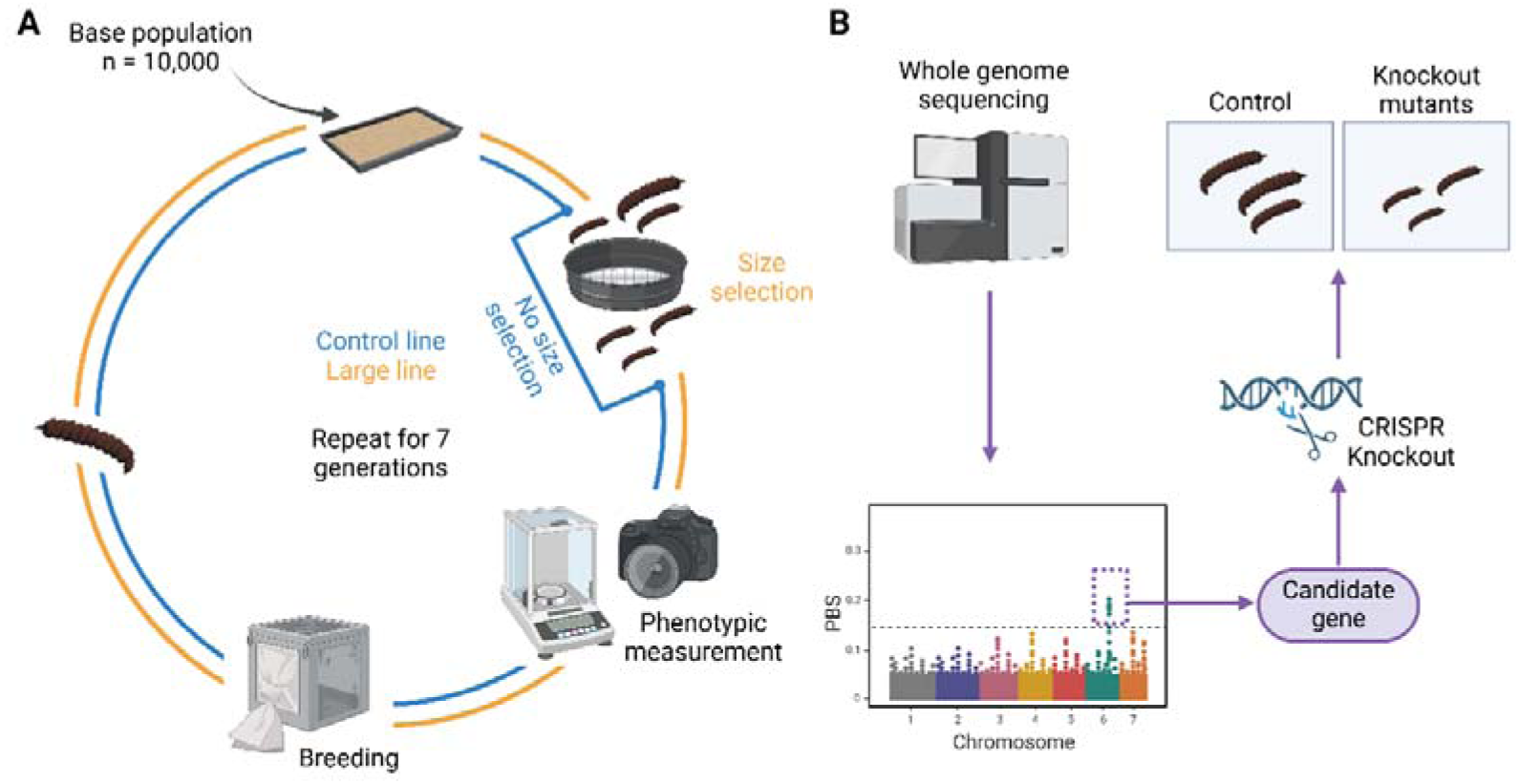
**A** pipeline for experimental evolution of pupal body size. Three replicate pairs of experiments were established using a starting population of 10,000 individuals (base population). For each replicate, one line underwent no size selection (control population) and another with selection for large pupal size (large selected population). Approximately 2,000 individuals were used to establish each line. Size phenotypes were estimated using image analysis. Larvae were reared to pupation in trays at a density of 0.5 larvae/cm^2^. **B** Pipeline for sequencing and analysis. From each of the three populations, 96 individuals were sequenced at low coverage (∼3x), and the Population Branch Statistic (PBS) was used to measure the differentiation of the large population relative to base and control, calculated in 20 kb non-overlapping sliding windows. Windows with PBS value higher than four times standard deviation were identified as “PBS peak regions”. A candidate gene was identified from genome scans and tested using a CRISPR knockout experiment. Figure generated using BioRender.

## Results

### Repeatable phenotypic response to selection for increased body size

Using a sieving technique to select the largest individuals, we obtained experimentally evolved populations which showed a consistent increase in body size across three replicates (replicate 1, +12.2%; replicate 2, +16.7%, and replicate 3, +16.5%). Pupae in the large selected line were significantly larger (+15% on average) than those of the control lines at generation seven (Figure 2A-C; Supplementary Table 1; Supplementary Table 2). Pupal size has a strong positive correlation with pupal weight (*R* = 0.85 – 0.9; *p* < 0.001) across all lines, indicating that body size (cm^2^) is also a good measure of weight (Supplementary Figure 1).

**Figure 2.**
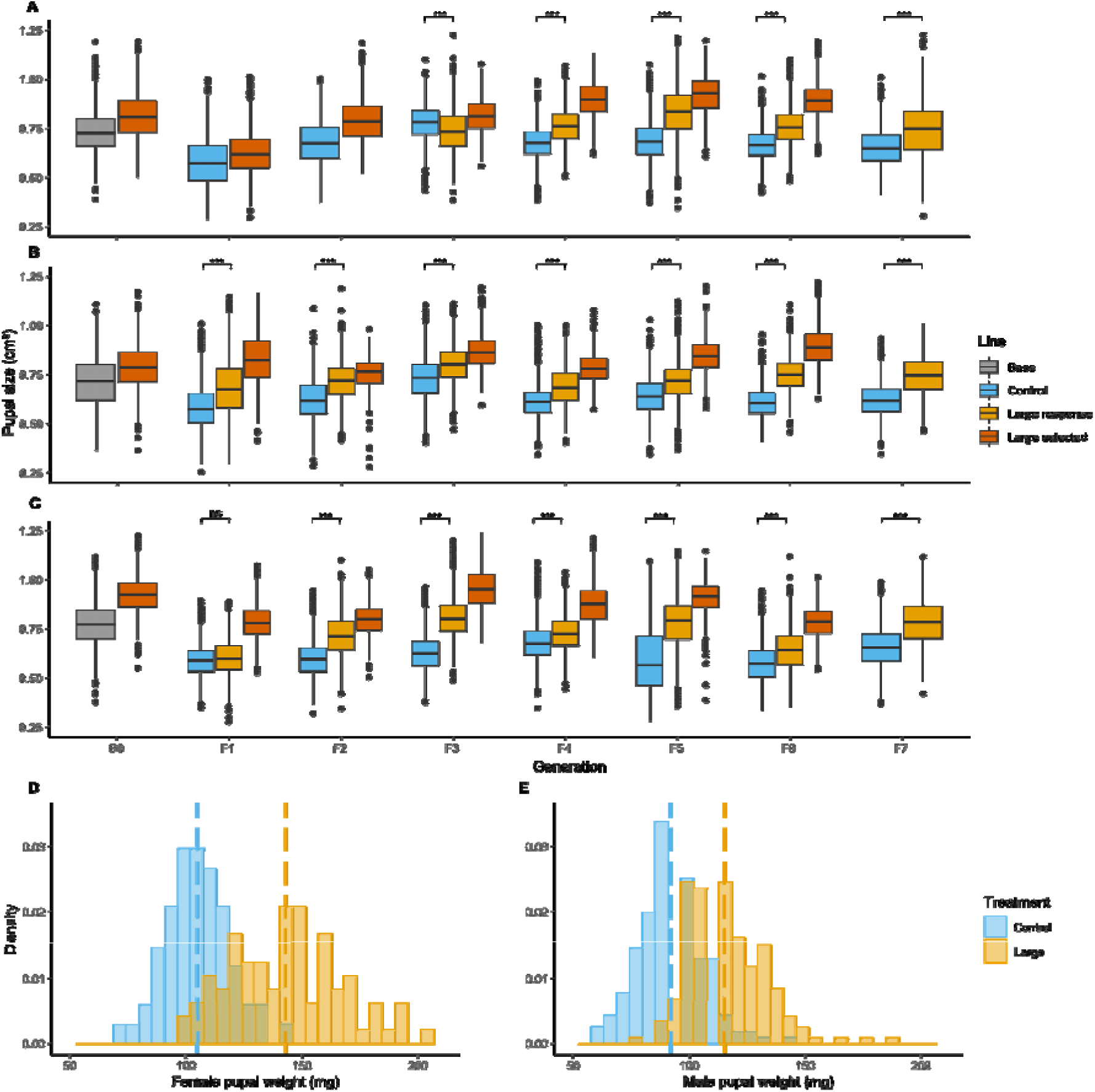
Evolutionary response to selection for large pupal body size replicated across three experiments (**A**-**C**) for seven generations. For each generation the pupal area (cm^2^) of size-selected individuals (selected) and that of their resulting offspring (response) is shown, as well as the base and control for every generation. Statistical significance was measured using a linear model and pairwise comparisons are presented using the Tukey post-hoc tests on the line*generation interaction (ns > 0.05, * < 0.05, ** < 0.01, *** < 0.001). Distribution of pupal weights of male (**D**) and female (**E**) individuals split by sex in the control and large-selected lines. Mean pupal weights from all replicates at generation seven are indicated with a vertical dashed line.

The response to selection was highly repeatable both across replicates and through generations (Figure 2 and Supplementary Table 2). The direction of change in body size was not always consistent between generations, likely due to stochastic variation in rearing conditions. Nonetheless, pupae were larger in the selected line as compared to the controls in every generation apart from the F3 of the first replicate (Figure 1A), in which selected individuals were actually smaller. In the F1 generation of the third replicate, selected individuals were larger as expected but the difference was not significant (Tukey-adjusted: *p* = 0.052; Figure 1C). Both selection line (LMM: *F*_1_,_77,219_ = 11,569, *p* < 0.001) and generation (LMM: *F*_6_,_77,219_ = 1,216, *p* < 0.001) were significant predictors of size across replicates. The significant line*generation interaction term (LMM: *F*_6_,_77,219_ = 141, *p* < 0.001) further indicates an increase in size in selected lines compared to controls across the generations. Only 3% of observed size variation was explained by replicate as a random effect, demonstrating high repeatability in the selected phenotype. Pupal body size of the response lines increased from 0.628 ± 0.12 cm^2^ in the F1 to 0.757 ± 0.12 cm^2^ in the F7 generation (Supplementary Figure 3). Body size showed a rapid and dramatic increase in the first three generations, with a less rapid change in subsequent generations (Figure 2; Supplementary Figure 3). From generation three onwards, the selected lines showed sizes (F7: 0.757 cm^2^) significantly greater than those observed in the original base population (G0: 0.732 cm^2^). Using the Breeders Equation we estimated that the heritability (*h^2^*) of pupal body size was 0.36 across our experiment, with values across replicates and generations ranging between 0.19-0.62. In summary, selection was effective in generating larger flies in a highly repeatable manner across replicates.

### Sex differences in selected lines

As expected^22^, female pupae were larger than males in both the control (12%) and large-selected lines (19%) (Figure 2D-E). Furthermore, pupal weight increased in both males (control: 92.21 ± 13.2 mg; selected: 115.39 ± 15.8 mg) and females (control: 104.97 ± 13.9 mg; selected: 143.12 ± 24.6 mg) after seven generations of selection (Supplementary Table 3), indicating that selection was effective on both sexes. Sex (*F*_1,556_ = 116.6, *p* < 0.001), line (*F*_1,556_ = 116.6, *p* < 0.001) and a sex*line interaction (*F*_1,556_ = 11, *p* < 0.001) were the strongest predictors of pupal weight (Supplementary Table 4).

Selection for large pupae will inevitably lead to a female-biased founding population for the selected lines, and we also saw a non-significant trend towards an increase in the proportion of females in the selected lines (9% more females across replicates, *X*^2^ = 1.35, *p* = 0.25), suggesting that we may have inadvertently selected for a female-biased primary sex ratio. This increase in females was found across the first two replicates but not the third (Supplementary Table 3). Ovary weights were also not significantly different between lines (*t* (106) = -1.45, *p* = 0.15), despite being 2% larger on average (Supplementary Table 3). A weak correlation between female weight and ovary size suggests that larger females do not consistently contain larger ovaries (*R* = 0.14, *p* = 0.18; Supplementary Figure 4).

### Little evidence for life history trade-offs

Genetic change in a trait such as body size alters resource investment which can lead to fitness trade-offs. We investigated whether selection for increased body size imposed trade-offs across other traits. First, size related traits increased in selected lines across all life stages. On average, larvae were 16% heavier in selected lines (Supplementary Table 5). Fixed effects of line (*F*_1,9_ = 46.1, *p* < 0.001) and day (*F*_15,7558_ = 1595, *p* < 0.001) were strong predictors of increased larval weight (Supplementary Table 6). Replicated experiments as a random effect alone explained 17% of variation (Supplementary Table 6). Additionally, random batch effects across rearing trays within each replicated experiment explained just 3% of variation.

Not only did larvae grow larger in the selected line, but they also contained significantly more protein (LMM, *F*_1,54_ = 15.26, *p* < 0.001; +13.7%) (Figure 3C; Supplementary Table 3). Random effects of replicate and rearing tray explained 42% and 19% of variation in protein content respectively (Supplementary Table 7).

**Figure 3.**
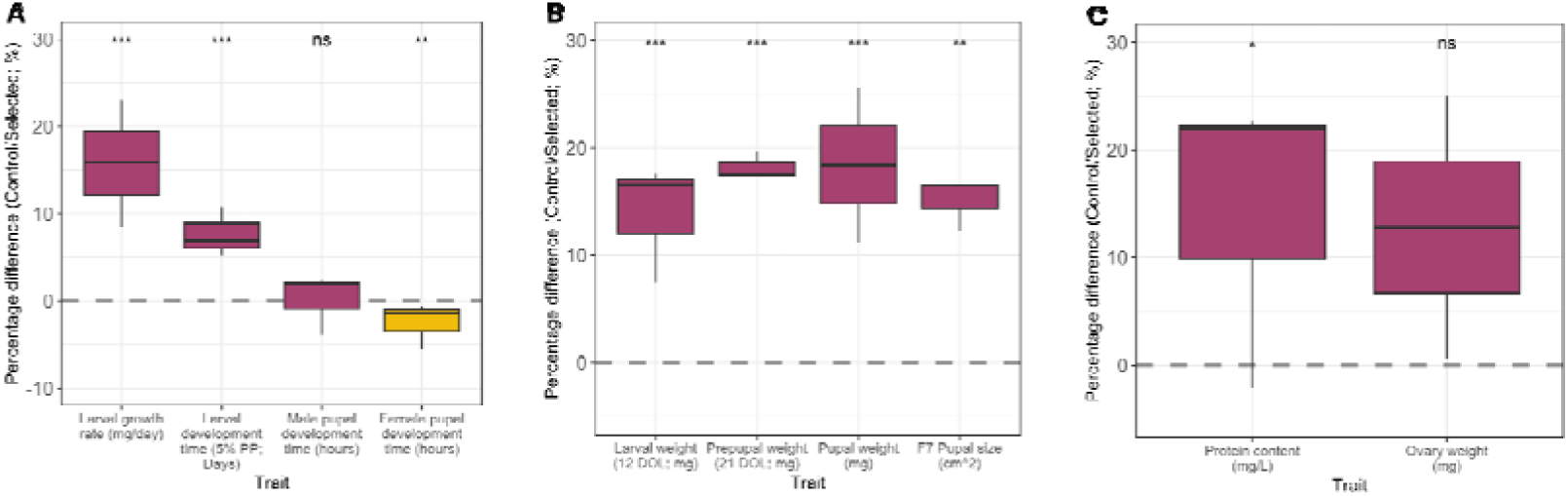
Percentage change in growth (**A**) and weight (**B**), larval protein content and female ovary weight (**C**) between selected and control populations. Traits with an increase in selected strains compared to control strains are shown in purple, and traits showing a decrease in orange. Results from comparisons using t-test and Wilcoxon tests for all traits are shown (ns > 0.05, * < 0.05, ** < 0.01, *** < 0.001).

Development time is another trait likely to show a trade-off with size. Large individuals might be expected to take longer to develop. In particular, pupae are typically harvested at the point at which the population reaches ≥ 95% larva/≤ 5% prepupa^23^. However, in replicate 1 and replicate 2 the selected lines reached the harvest stage three and two days faster respectively than controls. Replicate 3 showed no difference between lines (Supplementary Table 8). On average across all replicates, the harvest threshold was reached on day 13 in the large-selected line (89.08%) and on day 14 in the control line (93.63%; Supplementary Table 8). The proportion of larvae on day 13 was significantly lower in the large-selected line than the control, indicating faster larval development (*X*^2^ = 5.39, *p* < 0.05). However, differences disappeared by day 16 (*X*^2^ = 0.32, *p* = 0.57) ( Supplementary Table 8).

Nonetheless, larger individuals showed a longer pupal, but not larval, development time in both control (*R* = 0.19, *p <* 0.01) and selected (*R* = 0.28, *p* = < 0.001) lines (Supplementary Figure 5). However, when comparing selected and control lines, this difference was only significant for females, who invest substantially more in body size than males, with a pupal development time 6% longer in the selected lines compared to controls (*p* < 0.01) (Figure 3A, Supplementary Figure 4H). Male pupal development time did not differ significantly between controls and selected lines (*p* = 0.43) (Figure 3A, Supplementary Figure 4H).

### Polygenic and non-repeatable genomic landscape after body size selection

To study the genetic response to selection, we shotgun sequenced the whole genome of 288 individuals from base, control and large selected populations (32 per population) across the three replicates. We first tested for the influence of inbreeding and found that inbreeding coefficients per individual showed no significant change in any pairwise comparison in all three replicates (Supplementary Figure 6). However, the large selected line in every replicate has a decreased population-wide average π value compared to base (-0.0006, -0.003 and -0.0017 in 1, 2 and 3, respectively) and control (-0.0005, -0.0022 and -0.0009 in 1, 2 and 3, respectively) lines, showing an overall impact of selection on nucleotide diversity (Supplementary Table 10).

PCA analysis showed genetic differentiation from the base population in both control and selected lines (Supplementary Figure 7), but that selected lines tended to show a larger distance from the base, with replicate 2 quite distinct from the other two replicates. There was no obvious clustering between female and male individuals in the PCA (Supplementary Figure 7A).

Unselected lines were designed to evaluate the effects of genetic drift. *F*_ST_ was highly heterogeneous in control-base comparisons but was much lower as compared to large-base (Supplementary Figure 8B, D, F). This demonstrates that although genetic drift likely does alter the genomic landscape during experimental evolution, the influence of selection was still highly evident in selected lines (Supplementary Figure 8A, C, E).

We next used the Population Branch Statistic (PBS) to search for signatures of divergence in the selected line. The PBS is an *F*_ST_ based statistic designed for a three-population comparison^24^, with the control-base comparison effectively controlling for the effects of drift during selection. Using a conservative significance threshold of four times the standard deviation (4SD)^25^, the PBS statistic showed a highly heterogeneous genomic landscape with multiple significant peaks on different chromosomes in each replicate (red solid lines) (Figure 4A). The genomic landscape in replicate 2 was more heterogeneous than those in the other two replicates.

**Figure 4.**
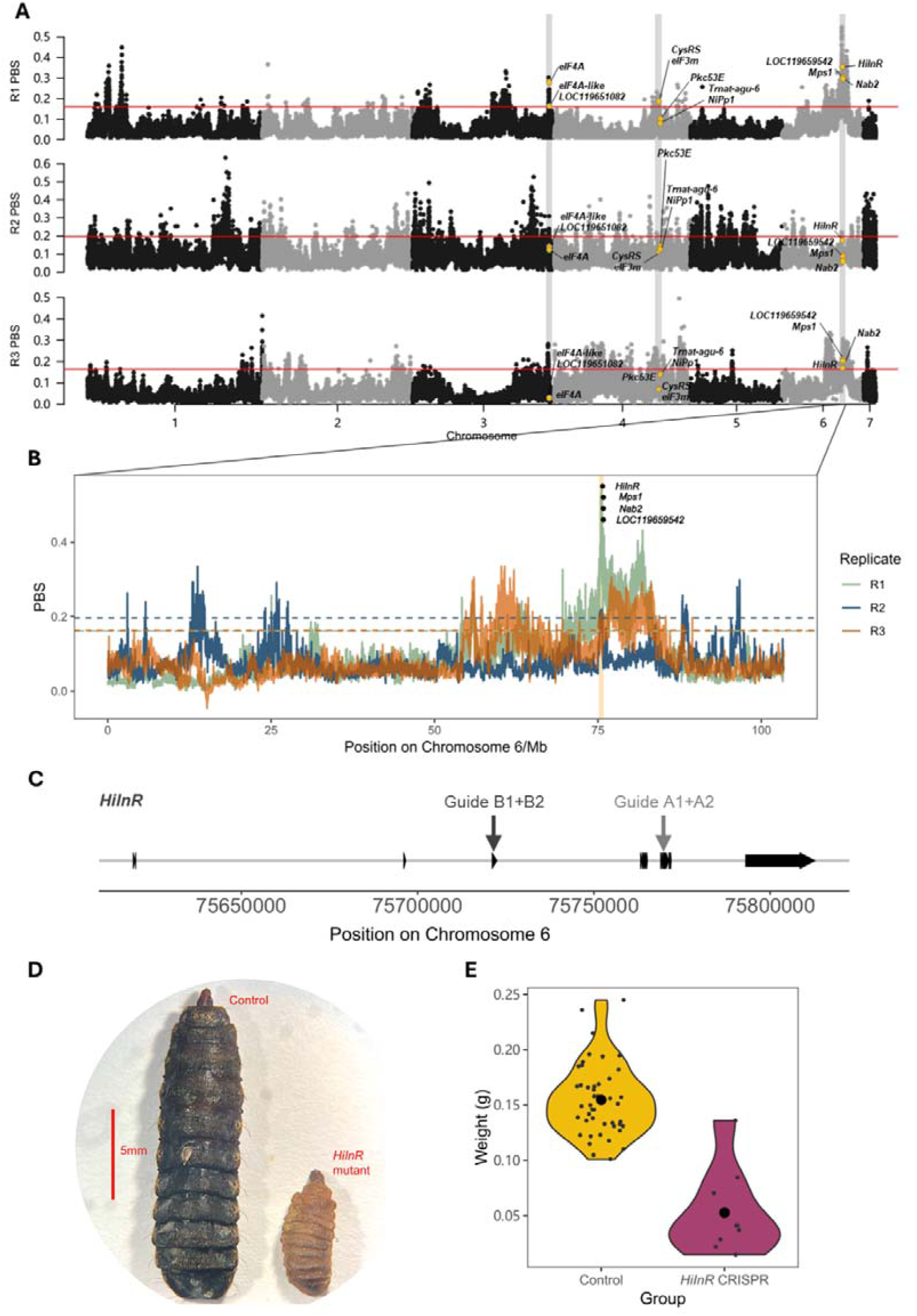
**A** PBS landscape of the large selected line compared to base and control populations shown for each replicate. The PBS statistic uses two populations as a baseline (in this case, control and base) and identifies genome regions that represent *F*_ST_ outliers in the selected line as compared the other two populations. We used an arbitrary threshold of four times the standard deviation of PBS values (‘4SD’ hereafter)^25^. Adjacent chromosomes are distinguished with black and grey colours. The solid red line in each landscape represents the 4SD of PBS values in the same replicate. The 12 candidate genes found in shared regions are highlighted in yellow. **B** PBS landscape at the overlapping peak on Chromosome 6. The 75160000-75840000 bp peak region is highlighted in the rectangle. Threshold values of 4SD shown by horizontal dashed lines. Candidate genes within the peak region are mapped to the midpoint between their start and end positions. **C** Simplified illustration of the *HiInR* gene structure, with exons represented as black arrows and introns as grey lines, and two pairs of CRISPR sgRNA guides - A1, A2 targeting exon 16 and B1, B2 targeting exon 4. **D** Body size comparison between a *HiInR* mosaic mutant and a control individual at day 28 post CRISPR injection. **E** Weight of *HiInR* mosaic mutants and control group at 28 days post CRISPR injection (unpaired Welch’s t-test, *p* **< 0.001**).

Sliding windows with PBS values above the threshold were extracted and a 100 kb buffer zone was added both upstream and downstream to account for gaps between sliding windows in a potentially continuous genome region^25^. After combining windows in this way, 120, 301 and 157 peak regions were identified in replicate 1, 2 and 3, respectively, indicating a highly polygenic response to selection. These high divergence windows also showed evidence for an excess of high-frequency alleles in the selected population as compared to base and controls, providing further evidence of selection in large lines (Supplementary Figure 9). Nonetheless, repeatability was low between replicates, with only four regions shared across all three replicates, located on three chromosomes (Chromosomes 3, 4 and 6; Figure 4; Supplementary Table 12).

### Candidate genes involved in pupal body size difference

We next examined the overlapping PBS outlier windows for candidate genes and found only twelve genes within the four overlapping outlier regions (Figure 4A). In the original 20 kb sliding windows without the 100kb buffer zones, only five sliding windows were shared across all three replicates (Supplementary Figure 10), all located within the Chromosome 6 75160000-75840000 bp peak which is also the longest peak, spanning 0.68 Mb in length in all replicates. Four genes (*LOC119659542*, *Nab2*, *Mps1*, *InR*) were found within this region. Among them, *Nab2* is the homolog of the human *ZC3H14* gene and has been shown to be relevant to brain development^26^, locomotion and alternative splicing^27^ in *Drosophila*. *Mps1* is involved in meiosis^28^ and chromosome segregation^29^ in *Drosophila*.

Within the Chromosome 6 peak, the gene *HiInR* encodes the insulin-like receptor and is part of the canonical growth regulation insulin signalling pathway^30,31^, making this a strong candidate for a functional role in size determination. We therefore conducted a CRISPR knockout experiment to generate *HiInR* mutants and confirm the expected role of this gene in regulation of larval growth. The *HiInR* gene was targeted using two pairs of CRISPR sgRNA guides (Figure 4C), but only guide pair A successfully generated mutants. Compared to the control group, *HiInR* mutants confirmed by genotyping have smaller larval body weight (Figure 4C, Supplementary Figure 11) (unpaired Welch t-test, df = 10.23, *p* < 0.001, Figure 4D). A delay in larval development was also observed in the overall *HiInR* CRISPR treatment group, represented by the smaller proportion of fully developed pupae and large number of larvae compared to the control group (*X*^2^, df = 2, *p* = 0.003, Supplementary Figure 12).

The *HiInR* gene is therefore a strong candidate for controlling some of the body size variation seen between control and selected lines. We found a single non-synonymous site which increased in frequency from the control to large selected line in two of the three replicates, replicate 1 (Fisher’s exact test, *p* = 0.007) and 3 (Fisher’s exact test, *p* = 0.013). This substitution caused an amino acid change from ‘T’ (Threonine) to ‘I’ (Isoleucine) at position 1,328 in the protein sequence and was the only consistent change in amino acid sequence between control and large selected populations found in more than one replicate. This therefore represents a candidate functional change but cannot explain the role of this gene in all three replicates.

## Discussion

Our replicated selection regime involved measuring 98,972 pupae, whole genome sequencing 288 individuals and demonstrated a strikingly repeatable phenotypic response to selection. Selected pupae grew 15% larger across replicates (Supplementary Table 1), a similar response to size selection as seen in a previous experiment over considerably more generations (22% in a previous study by Facchini et al.^20^). Nonetheless, as predicted for a highly polygenic trait, the genomic response was very different in each replicate, with between 120-301 selected regions identified in each replicate, but only 12 shared across all three. Despite the general lack of overlap between replicates, we identified one gene, the insulin receptor *HiInR* that was consistently under selection and whose function in growth regulation was confirmed using a CRISPR knockout experiment.

Life history traits such as size, development time, fecundity, longevity, and immunity, are constrained by the availability of resources and often show trade-offs^32^. In contrast to a previously demonstrated trade-off between weight and development time in BSF^33^, we here found a surprising lack of trade-offs, with selected larvae generally growing faster and having increased protein content and ovary weight. Only female pupal development time showed a trade-off, with larger female pupae developing more slowly. Interestingly, in *Drosophila*, trade-offs between longevity and body size are similarly strain specific^11^. Considering the genetic diversity in *H. illucens* it is likely that other strains may have evolved alternative resource investment strategies^34–36^, so these results may turn out to be dependent on genetic background.

Polygenic traits are expected to show high redundancy between loci in the response to selection. In mammals, body size is typically encoded by hundreds of genes^37^. However, there are also many large effect variants for body size. For example, canine insulin growth factor-1 (*IGF1*) is a key contributor to small body size^38^. In insects, evolve and resequence experiments in *Drosophila melanogaster* have shown a highly polygenic architecture with hundreds of genes controlling body size^39^. Similarly, our study shows that in all three replicates, a large number of sliding windows could be found in PBS peaks that exceeded threshold values. However, most PBS peaks appear in only one replicate, and only 12 genes were found within shared peaks across all three replicates. This indicates not only polygenicity, but also low repeatability in the genetic architecture of body size. This lack of repeatability across replicates implies genetic redundancy, such that different gene combinations and interactions can result in similar trait outcomes^5,6^. Body size is a complex trait that can reflect a result of resource acquisition and allocation during growth. The genes showing higher differentiation may therefore be relevant to different aspects of the phenotype that ultimately influence growth.

On Chromosome 6, the *HiInR* gene spans half of the shared peak region and is predicted to encode the insulin-like receptor, a protein that plays a significant role in growth regulation, metabolism and lifespan in many organisms^40–42^. Studies in *Drosophila* show that this signalling pathway is evolutionarily conserved and regulates individual size^43^. To confirm this function, we demonstrated both a delay in development and a decrease in body size in *HiInR* knockout mutants (Figure 4D&E). We identified a coding sequence change, located in an extracellular domain of the protein, that shows a significant frequency change in two of the three replicate lines and is therefore a candidate functional change. This amino acid is within the second Fibronectin Type-III (FN3) domain of the protein and is only five amino acids away from a predicted cytokine receptor motif within the FN3 domain. Nonetheless, this mutation cannot explain the same divergence peak around this gene in replicate 2, and other putatively regulatory mutations in non-coding sequence at linked sites could also be involved in the body size difference. However, despite low repeatability in loci reflected by the genomic landscapes, our evolve and resequence approach proves effective in identifying a strong candidate gene for this highly polygenic trait.

We designed our mass selection experiment to maintain large population sizes throughout the selection regime to minimise the effects of drift, and across replicates our experiment had an average population size of 1,703 individuals each generation. For all three replicates this resulted in no significant differences in the inbreeding coefficient in any pairwise comparison between base, control and selected populations (Supplementary Figure 6).

One previous study in black soldier fly attributed a loss of genetic diversity to drift in selected populations after just three generations of selection^21^. Drift is also likely to generate false positives in *F*_ST_ outlier analyses^44,45^, and to account for this we designed the genetic analysis with a control population for each replicate, and using the population branch statistic which accounts for change in the control population when identifying outlier loci^25,46^. Nonetheless, the higher number of outliers in our replicate 2 (301 as compared to 120 and 157 in the other two replicates) might indicate false positives due to drift. However, given the lack of significant inbreeding, and the much higher *F*_ST_ peaks in large vs base as compared to control vs base comparisons, it seems likely that most of the outlier peaks identified here under our stringent significance threshold are driven by selection.

## Conclusion

Our results support the expectations of a highly polygenic architecture, with many loci and low genetic repeatability, despite similarity in phenotype between replicates^1^. This indicates high genetic redundancy, with many different genetic routes that can generate a similar response to selection. This also implies large amounts of standing variation, and multiple genotypic solutions to optimising critical traits, which bodes well for selection to improve this species for industrial applications^7^. Nonetheless, despite little repeatability between replicates, we have identified a single gene involved in the canonical insulin signalling pathway that showed a consistent response to selection, with a coding change that represents a putative causal mutation selected in two of the three replicate lines. Overall, our results demonstrate the power of replication in experimental evolution studies to investigate the genetic basis of polygenic phenotypic traits.

## Materials and Methods

### Establishing experimentally evolved populations of *Hermetia illucens*

Source populations of *H. illucens* were obtained from Better Origin (Entomics Biosystems Limited, UK) and were maintained in the Zoology sub-department for animal behaviour at University of Cambridge (Cambridge, UK). Previous genome sequencing analysis indicates the source population (CAM006606), belongs to lineage-*a* of the *H. illucens* phylogeny (named as Industry-strain-B, in Generalovic et al. 2023). This population was maintained under industry conditions on a diet of poultry feed, with a feeding ratio ranging from 1.93 to 2.68 g per larvae (g/larvae). To minimise the effects of genetic drift and false positive rates on genomic signatures of selection, a selection program was designed based on previous studies^48^. Three independent replicates were established in March (replicate 1), May (replicate 2) and October of 2021 (replicate 3), using the core breeding stock from Better Origin operations. To establish each line per replicate, 10,000 individuals were obtained at each of the three stated time-points and an approximate effective population size of 2,000 random individuals were used to establish each line. Across selected and control lines there was an average population size of 1,703 individuals per generation (supplementary table 1). Establishing an internal control for each replicate and maintaining large population sizes was intended to both control for and minimise the effects of genetic drift.

### Imposing artificial selection pressures on established *Hermetia illucens* lines

Artificial selection was performed for increased pupal body size using a semi-automated, low-cost and high-throughput pipeline (supplementary figure 1). Body size was measured from a random sample of approximately 2,000 individuals from the source population. Individuals were imaged using a smartphone camera (Samsung S10, 1.5x fixed magnification) and images processed in ImageJ^49^ using a custom script^50^ to obtain mean “area” of pupal bodies. Hereafter “area” is referred to as “body size”, unless explicitly stated. A manual scale was set for every batch of images captured. This initial random sampling of pupae (G0, referred to as “base population” hereafter) was then used to establish the control line. The selection line was established by passing the remaining pupa (*ca*. 8,000) from the source population through a selection treatment using a mechanical sieve (Henan Remont I/E, China) with a custom mesh (4 mm^2^ mesh size). Preliminary optimisation indicated that 4 mm^2^ was efficient at separating the largest pupae at an adequate population size (data not shown). Pupa were added as input to the size selection treatment and mechanically shaken for two intervals of 1 minute allowing smaller individuals to pass through the mesh, remaining individuals were collected for imaging. Once imaged, every batch of pupae in both lines was weighed and recorded. Size selection was maintained for seven generations in the selected line and the response to selection was recorded every generation (F1-F7) except for the F1 and F2 of the first replicate. Pupal batches from each image were grouped independently for each line and mixed with approximately 200 g (< 5 cm in depth) of woodchips within an open topped 7 L tupperware container (Sistema, New Zealand). Pupal-substrate mixtures were then moved to a breeding cage (40*40*60 cm; Qiansha, China) within an environmentally controlled greenhouse.

### Experimentally evolved line maintenance

The experimental control and selected lines were maintained under identical environmental conditions for the duration of the experiment. Egg clutches of the F1s were collected synchronously from the breeding cages by replacing egg traps 24-hours prior to egg harvest, ensuring family age was fixed by day. Egg traps (four corrugated correx sheets, 0.3*4*8 cm) were placed over an oviposition attractant, comprised of ca. 500 g one-week-old stock feed, within a ventilated 1.9 L Tupperware container (Sistema, New Zealand). Egg mass was weighed, recorded and setup in a hatching tray. The open-topped hatching tray (Euronorm tray, 20*40*60 cm) was filled with 1 kg of stock feed encircled by 500 g of wheat bran (Burnhills, UK), and egg traps were placed in the centre. Hatch trays were maintained within a temperature-controlled environment. Each tray was enclosed within a ventilated net for three days to promote a humid environment and aid hatchability. On day three, ventilated nets were removed, and the larva-substrate mixture was allowed to dry. On day five, larvae formed aggregated pools amongst the frass that allowed frass-free collection of larvae for density correction. Density correction was performed on day five by taking a minimum of three 0.5 -1 g aliquots of larvae and counting number per sample. Mean larval weight per line was used to extrapolate required mass for an approximate density of 5,000 neonate larvae per tray. A maximum number of trays were seeded in each line depending on available neonate quantity reaching population sizes between 5,000 – 20,000 individuals. Seeding was performed into fresh rearing trays (Euronorm tray, 20*40*60 cm) with 1 kg stock-feed surrounded by 500 g dry wheat bran. Larval rearing occurred at an approximate density of 0.5 L/ cm^2^ and trays were stacked horizontally on a rack enabling air flow. Feeding occurred on a batch schedule every three days post density correction equating to a total of 6.5 kg (including hatch tray) feed input averaging 1.3 g/larvae. Trays were arranged alternating between control and selected lines and were rotated vertically after every feeding period. On the third day after the last feed was added, 500 g of wheat bran was mixed into each tray to dry out excess moisture and promote pupation. Pupae were subsequently isolated from the frass using a 2 mm^2^ aperture attachment sieve for the mechanical sieve (Henan Remont I/E, China). This did not remove any pupae from either of the lines. Isolated pupae from each line were then subjected to their line specific treatment. Control lines were imaged while selected lines were imaged to measure the response to selection, size-selected, and re-imaged to obtain size phenotypes. Both lines were then established in breeding cages as previously described. This program was repeated until generation seven. After generation seven, both lines were maintained without the selection regime and fed *ad libitum* for future work.

### Phenotypic assessment and estimates of heritability

Pupal body size phenotypes were measured using the image analysis pipeline as described above. Artefacts were detected in several images caused by frass particles (supplementary figure 2). Manual inspection of these images revealed the contaminants as outliers in pupal body size. These outliers, which measured below 0.2 cm^2^ and above 1.5 cm^2^, were removed.

After seven generations, more extensive phenotypic data was collected. Generation seven egg masses were collected in a synchronised manner, established within neonate hatching trays and density corrected in parallel as previously described. Daily mean larval weights were obtained from days five to 20 by measuring a maximum of 25 individual larva per tray in each line. This allowed daily weights and an average growth rate to be determined. Within-tray larval, prepupal and pupal development dynamics were measured from day five to 25 by acquiring three *ca*. 300 g samples of *H. illucens*-frass mixture and estimating the average proportion of each life stage present in each tray over the course of the experiment. Prior to daily measurements all trays were mixed thoroughly by hand to promote a homogenous distribution of sampled phenotypes due to larval aggregation and prepupae “self-harvest” behaviours.

Protein content was generated as average proximate body composition of three lyophilized and homogenised larvae sampled on day 12. Prepared samples were subjected to Carbon-Nitrogen-Hydrogen (CNH) combustion analysis (CE-440 Elemental Analyzer, Exeter Analytical) at the University of Cambridge Chemistry department. Proximate protein composition was calculated using a species-specific Nitrogen conversion factor of 4.76^51^. Collection of eight-to-12 pooled samples were performed for each experimental line across trays within lines.

Pupal development rates were measured to the nearest observed hour to identify differences between size and development within sexes. During daily tray investigations, pupae were collected and isolated into individual *Drosophila* rearing vials, with time and date recorded. Isolated pupae were checked frequently between the hours of 08:00 and 18:00 over ten days for adult emergence. Pupa that failed to emerge past ten days were deemed as fatalities. Total time for adult emergence was recorded as the maximum pupal development time. Freshly emerged adults were sexed, and a population sex ratio was inferred from the sampled emerged adults as a representative of the population. Ovary weights were collected by dissection of three-day old adult females previously isolated, weighed as an ovary pair and expressed as total ovary weight (not corrected for adult size).

Pupal size was recorded using the image analysis pipeline as previously described. All weight-based measures were obtained using a Mettler AE 163 (Mettler-Toledo, UK; sensitive to four decimal places). Size relationships between pupal width, length and weight were performed manually using a digital calliper (Machine-DRO, UK). At the base population (G0) and every subsequent generation (F1-F7) post-establishment, a sample of 100 larvae (sampled on day-12) and approximately 60 adults at a 50:50 male to female ratio were collected from both lines and stored in > 96% ethanol at -20 °C for future genomic analysis.

Estimates of narrow-sense heritability (*h^2^*) were recorded using the breeder’s equation^52^, R = *h^2^S*. The selection differential (*S*) was generated as the difference in mean trait values between the selected and control populations. Response to selection (*R*) was measured as the difference in mean size between generations in the selected line.

### Statistical analysis of phenotypes

All statistical analysis was performed in Rstudio v1.2.5 (RStudio Team 2019). Data visualisation was generated using the ggplot2 v3.3.6 R package^53^. Additional packages used include dplyr v1.0.5^54^, forcats v0.5.1^55^, Rmisc v1.5.1^56^, ggside v0.1.3^57^, lmerTest v3.1-3^58^ and emmeans v1.5.4^59^. A linear model (LM) using the control and the large-selected response pupal body size data for each replicate was used to measure the main effects of line, generation and the interaction driving the difference in size between lines. Post-hoc analysis using the Tukey correction method was performed on significant fixed effects. In addition, a mixed-effect linear model (LMM) was also performed across all replicates in a similar manner but incorporating (1| replicate) as a random effect. Assumptions of normality and homoscedasticity were assessed visually due to the large dataset (*n* = 77,234 datapoints).

Pairwise comparisons were performed for several phenotypes across all experiments at generation seven. A Welch two-sample T-test was performed for pupal size, larval protein content and ovary weight traits at generation seven. Unpaired two-sample Wilcoxon signed ranks test were performed for daily larval growth rates, fixed life stage weights (larval, prepupal and pupal), larval development and pupal development rate traits.

Each trait was then further explored using full statistical models to explore effect interactions and random variables. A LMM was performed to test for differences in daily larval weight over the growth period (fixed effects: line, day and line*day; random: replicate, tray:replicate). Post-hoc analysis was performed using Tukeys multiple testing adjustment. A sample of larval (day-12), prepupal (day-21) and pupal (day-18) weights at fixed days were sampled and differences were compared between lines using unpaired two-sample Wilcoxon signed ranks tests. Protein content data was log transformed and analysed using a LMM (fixed: line; random: replicate & tray:replicate). Larval development was measured within rearing trays and assessed using a two-proportion Z-test at each day. Pupal development time was first assessed by grouping sexes and comparing between lines using an unpaired two-sample Wilcoxon signed ranks test. A sample of pupae were also weighed and isolated until adult emergence to determine sex. Sexed pupal weight was analysed using an LMM, post-square root data transformation, to explore the effect of line, sex, and the interaction of the two (fixed: line, sex & line*sex; random: replicate & tray:replicate). Post-hoc analysis was performed using Tukeys multiple testing adjustment. Resulting adult sexes were counted as a proportion of females to males. Female proportion was analysed using a two-proportion Z-test. An unpaired two sample t-test was performed for each independent replicate when testing differences in ovary weight between lines post square-root transformation. All correlation analysis was performed using a linear regression model. Parametric model assumptions were assessed for normality with Shapiro-Wilk and equality of variances *F*-tests or visually. Data failing to meet assumptions of normality and homoscedasticity were analysed using non-parametric tests as described.

### Sample collection, whole genome sequencing and data pre-processing

Random samples from the 7^th^ generation of the control and selected lines and from the base population before selection were collected, covering two sexes and three replicates to generate a dataset of 288 individuals (Supplementary Table 9). All samples were stored in ethanol at –20 °C until sequencing.

Sample DNA was extracted using a bead-based protocol^60^. Sequencing libraries were made using homemade Tn5 for fragmentation and then barcoded and amplified with nextera primers^61^. Libraries of all individuals were pooled and whole-genome sequencing was performed on the Illumina platform with Novogene (Novogene, Cambridge, UK). Raw fastq data was quality controlled by fastQC 0.11.8^62^ both before and after adapters were removed with Trimmomatic 0.39^63^. Trimmed data were mapped to the chromosomal level reference genome^64^ using BWA 0.7.17^65^. Duplicated reads were removed using the MarkDuplicates function in Picard 2.9.2^66^.

All pre-processing steps were integrated using Snakemake pipeline^67^ to handle all individual sequences in batches. The deduplicated BAM files were then used for following analyses except for the principal components analysis, where variant calling was carried out and SNPs data was used.

For the principal components analysis, a variant calling step was performed using GATK 3.7.0^68^. Hard filter arguments to remove low quality sites were set as “MQ < 40.0, QD < 2.0, FS > 60.0, SOR > 3.0, MQRankSum < -12.5, ReadPosRankSum < -8.0, DP < 238.8, DP > 955.1” according to the GATK recommendation with adaptation based on the dataset.

### Principal components analysis

Principal components analysis (PCA) was performed using the genotyped variant data with PLINK^69^. 20 principal components were used in the analysis. Output eigenvec file was visualized in R using packages “ggplot2”^53^, “ggsci”^70^ and “ggpubr”^71^.

### Candidate gene selection

For each of the three replicates, three comparisons of *F*_ST_ (LARGE-BASE, LARGE-CTRL and CTRL-BASE) and population branch statistic (PBS) of the large selected line were calculated and normalised in non-overlapping 20kb sliding windows with the realSFS module of ANGSD^72^. As the PBS value makes use of all *F*_ST_ values among three populations, it takes into account the differentiation level that may be caused by drift (CTRL-BASE). A threshold of four times standard deviation of PBS values of the large selected population in each replicate was chosen for candidate genes identification. PBS landscape of each replicate was visualized with qqman^73^ R package.

As some peak regions are continuous and may consist of windows with PBS value both higher and lower than the threshold, we added a 100kb ‘buffer zone’ both upstream and downstream of each high divergence window to better join windows located within the same peak region. Windows with buffer zones added were then joined using a custom Python script to form a list of ‘peak regions’.

In order to get parts of the regions that are shared among all three replicates, the ‘intersect’ function implemented in the software BEDTools^74^ was used. First, the shared intersects between replicate 1 and replicate 2 were extracted using default arguments without adding new buffer zones. This generated a list of shared regions between the two replicates and their start and end positions, which were used as input for another round of intersect with replicate 3. Gene IDs were extracted from the annotation file (NCBI *Hermetia illucens* Annotation Release 100) generated by NCBI annotation pipeline based on a previously published reference genome^64^.

### Site frequency spectrum (SFS) estimation

In order to study the change in SFS pattern in selected regions of all populations, sliding windows of PBS values above the 4SD threshold were extracted separately and their SFS were estimated using ‘realSFS’ function in ANGSD. The output files of all separate windows were combined together using a custom script to form a single summary file of each population and each replicate. This was visualized with the basic barplot function in R.

### Inbreeding analysis

In order to investigate the possibility of inbreeding during the selection process, we calculated per-individual inbreeding coefficients (*F*) with ngsF^75^. To minimise the effect of variance in per-site sequenced individuals, a filter was used during the glf file generating process to exclude all sites with less than 30 individuals in each population. ngsF was run for each population with parameters “--init_values r --min_epsilon 1e-6” to calculate per-individual inbreeding coefficients.

### CRISPR gene knockout for *HiInR*

#### Guide design, assembly, in vitro test

CRISPR sgRNAs were designed in Geneious Prime (Version 2024.0.7. https://www.geneious.com) from a Chromosome 6 consensus sequence of 288 base, control and large selected line sequences. Two pairs of sgRNA guides targeting two different CDSs were selected for *HiInR* (Pair A: (5’-3’) TATGGCACTTCCGAAGGCGGTGG, (5’-3’) GCGATTTGAAAGGGTATCTACGG; Pair B: (5’-3’) GCTGAACCCTTGTCCCAGCG, (5’-3’) TAGCGACCAGACGAATACAG). sgRNA guides were ordered from IDT (Integrated DNA Technologies, Inc). Flanking primers were also designed in Geneious Prime to amplify the two target regions (Pair A: forward TCTGCTGCTTCTCTATTGTCTTT, reverse CGTCCCCTTTCTATAATAATCCGT, Pair B: forward TTATCGCTCCATGCACAAAATG, reverse TACAATATACTGCACCGGTCAC) and ordered from IDT.

#### Microinjections

Microinjections were performed as described in a previous study^76^ using quartz capillary needles (Sutter Instrument). Injection mixes were prepared using a mix of sgRNA pairs A and B (each at 125 ng/µl, giving a total concentration of 500 ng/µl), mixed with Cas9 at 500 ng/µl. Freshly laid eggs were collected within one hour of oviposition. A single clutch or a mix of egg clutches were split equally between a control group receiving a sham injection of Cas9 at 500 ng/µl without guides or injection mix as stated previously. Injections were performed within four hours of oviposition.

### CRISPR larval rearing and phenotyping

CRISPR injected groups and control groups were treated under identical rearing conditions. Eggs were incubated at 25 °C in petri dishes surrounded by moist tissue sprayed with 1% Calcium Propionate for two days. Embryos at two-days old were density corrected to the same population size and placed in agar-based food (1.5% agar, 2% potato flakes, 2% Wheat bran, 5% Glucose, 4% Whey, 0.2% Methionine, 0.2% Tyrosine, 0.1% Calcium Propionate, 0.74% egg powder, 83.1% non-sterile water, 0.1% silage mix (P*ediococcus pentosaceus, Lactobacillus plantarum* and *L. brevis* and 1% dried yeast), with 2-3g of wet feed allocated per larva, providing a large excess of food to allow larvae to feed *ad libitum* and ensure that food is not a growth limiting factor. Larvae were cleaned, corrected to equal densities for control and CRISPR groups and provided with 2-10g of poultry feed based food (66.6% Allen & Page Layers Pellets chicken feed, 33.3% bran, 0.1% *Lactobacillus*, mixed at 3:7 with water) to allow larvae to feed *ad libitum* at 11 days post injection. Larvae were cleaned, photographed, weighed and density corrected at days 15, 20 and 28 after injection, with 2-10g of poultry feed based diet provided per larva on day 15 and 20. Density correction was performed at regular intervals on day 11, 15 and 20 to account for larval mortality throughout development and maintain CRISPR and control groups at even larval densities. At day 20, larvae were moved to 4 L plastic containers with perforated lids (Sistema, New Zealand) with an additional 5g of wheat bran to promote pupation. On day 28, larvae were photographed, weighed and genotyped using the same method described above.

To compare the mean weight difference between *HiInR* mutants and control group, unpaired Welch’s T-tests were performed using the *rstatix*^77^ and *magrittr*^78^ R packages.

### CRISPR mutant screening

Whole larvae and head capsules were used to extract DNA from individuals using a one-step extraction protocol^79–81^. Unpurified DNA lysate was diluted to 1:10 then PCR amplified using primers for respective guide targeted regions and NEB OneTaq® Quick-Load® 2X Master Mix, following manufacturers recommended protocol. PCR products were bead purified and Sanger sequenced Genewiz, UK. Sequenced reads were aligned against a wildtype sequence from the control group in Geneious to screen for gene knockouts. The presence of multiple low read certainty chromatogram traces originating within or close to the CRISPR guide target that persist downstream, and were not present in wild-types, were indicators of CRISPR mediated deletions. Nine individuals were found with CRISPR guide pair A mediated deletions. Guide pair B did not mediate deletion in any individual in the experiment.

## Data availability

All raw sequences used in this study has been uploaded to ENA database (Project: PRJEB70887). Raw data of phenotypic measurements used in this study have been released on https://github.com/TGeneralovic/BSFEvolveandResequenceMS.

## Code availability

Scripts used in this study have been released on https://github.com/TGeneralovic/BSFEvolveandResequenceMS.

## Acknowledgements

This study is funded <u>by the Biotechnology and Biological Sciences Research Council (BB/M011194/1) NPIF studentship awarded to Tomas Generalovic</u>. We thank Better Origin for support in genomic sequencing and providing the original base populations for the experiments. Wenjun Zhou acknowledges support of China Scholarship Council and Cambridge Trust (Project ID: 202206380035).

## Author contributions

CDJ, TG, and MP conceived and designed the study. TG carried out the selection experiment with help from SL. IW prepared libraries for sequencing and data analysis of genome data was carried out by WZ. The paper was written by TG, WZ and CDJ, with input from all other authors.

