## Supplementary figures and tables for "Repeatable phenotypic but not genetic response to selection on body size in the black soldier fly"

**Supplementary information**

| **Supplementary Table 1.** Summary statistics for pupal body size averaged across the three replicate experiments. Area (cm^2^) measurements for body size in the control, response to selection (large-response) and selected (large-selected) populations are provided with standard deviations. Size was recorded for each replicate and line from the base population to the seventh generation (F1-F7). | | | | | | |
| --- | --- | --- | --- | --- | --- | --- |
| **Generation** | **Control** | | **Large-response** | | **Large-selected** | |
|  | ***N*** | **Area (cm^2^)** | ***N*** | **Area (cm^2^)** | ***N*** | **Area (cm^2^)** |
| F1 | 5,160 | 0.583 ± 0.106 | 2,853 | 0.628 ± 0.117 | 3,939 | 0.710 ± 0.137 |
| F2 | 5,570 | 0.631 ± 0.106 | 3,490 | 0.715 ± 0.107 | 3,383 | 0.784 ± 0.097 |
| F3 | 5,616 | 0.705 ± 0.118 | 6,297 | 0.785 ± 0.108 | 2,236 | 0.892 ± 0.114 |
| F4 | 6,505 | 0.657 ± 0.092 | 5,745 | 0.725 ± 0.096 | 4,168 | 0.850 ± 0.104 |
| F5 | 6,115 | 0.641 ± 0.129 | 5,819 | 0.776 ± 0.128 | 4,061 | 0.897 ± 0.091 |
| F6 | 5,627 | 0.614 ± 0.093 | 5,866 | 0.718 ± 0.110 | 3,189 | 0.857 ± 0.103 |
| F7 | 6,076 | 0.641 ± 0.092 | 6,166 | 0.757 ± 0.116 | 1,091 | 0.867 ± 0.094 |

| **Supplementary Table 2** Descriptive statistics on pupal body size phenotypes from the base population to subsequent F7 progeny in both the control and selected lines (L-). Population sizes every generation and mean pupal body size (cm^2^) is provided with standard deviations. Data presented by replicate and missing data is denoted by NA's. | | | | | | | | | | |
| --- | --- | --- | --- | --- | --- | --- | --- | --- | --- | --- |
| **Gene-ration** | **Line** | **Replicate 01** | | | **Replicate 02** | | | **Replicate 03** | | |
|  |  | ***N*** | **Size (cm2)** | **SD** | ***N*** | **Size (cm2)** | **SD** | ***N*** | **Size (cm2)** | **SD** |
| G0 | Base | 1,656 | 0.732 | 0.109 | 2,098 | 0.715 | 0.122 | 2,043 | 0.773 | 0.107 |
| F1 | Control | 1,670 | 0.575 | 0.121 | 886 | 0.615 | 0.129 | 1,989 | 0.589 | 0.082 |
| F1 | L-response | NA | NA | NA | 867 | 0.690 | 0.142 | 1,986 | 0.600 | 0.091 |
| F1 | L-selected | 2,146 | 0.807 | 0.111 | 1,800 | 0.788 | 0.108 | 1,992 | 0.923 | 0.089 |
| F2 | Control | 1,702 | 0.675 | 0.112 | 1,869 | 0.624 | 0.102 | 1,999 | 0.599 | 0.090 |
| F2 | L-response | NA | NA | NA | 1,497 | 0.715 | 0.105 | 1,993 | 0.716 | 0.108 |
| F2 | L-selected | 1,939 | 0.621 | 0.106 | 608 | 0.823 | 0.132 | 1,392 | 0.785 | 0.087 |
| F3 | Control | 1,535 | 0.776 | 0.095 | 2,058 | 0.729 | 0.111 | 2,023 | 0.627 | 0.093 |
| F3 | L-response | 1,755 | 0.737 | 0.112 | 2,460 | 0.801 | 0.099 | 2,082 | 0.807 | 0.102 |
| F3 | L-selected | 1,819 | 0.787 | 0.108 | 657 | 0.755 | 0.081 | 907 | 0.798 | 0.079 |
| F4 | Control | 2,022 | 0.677 | 0.085 | 1,983 | 0.610 | 0.081 | 2,500 | 0.679 | 0.093 |
| F4 | L-response | 1,717 | 0.761 | 0.089 | 2,037 | 0.691 | 0.097 | 1,991 | 0.728 | 0.089 |
| F4 | L-selected | 483 | 0.812 | 0.093 | 825 | 0.867 | 0.085 | 928 | 0.956 | 0.111 |
| F5 | Control | 2,035 | 0.685 | 0.100 | 2,136 | 0.643 | 0.096 | 1,944 | 0.593 | 0.167 |
| F5 | L-response | 2,010 | 0.832 | 0.127 | 1,819 | 0.714 | 0.096 | 1,990 | 0.776 | 0.129 |
| F5 | L-selected | 1,239 | 0.895 | 0.091 | 1,419 | 0.782 | 0.076 | 1,510 | 0.876 | 0.105 |
| F6 | Control | 1,997 | 0.666 | 0.078 | 1,482 | 0.604 | 0.077 | 2,148 | 0.574 | 0.094 |
| F6 | L-response | 2,192 | 0.759 | 0.095 | 1,686 | 0.752 | 0.085 | 1,988 | 0.643 | 0.106 |
| F6 | L-selected | 1,262 | 0.921 | 0.096 | 1,044 | 0.847 | 0.084 | 1,755 | 0.910 | 0.081 |
| F7 | Control | 1,910 | 0.652 | 0.091 | 2,238 | 0.620 | 0.083 | 1,928 | 0.655 | 0.099 |
| F7 | L-response | 2,035 | 0.743 | 0.133 | 2,109 | 0.744 | 0.094 | 2,022 | 0.785 | 0.115 |
| F7 | L-selected | 929 | 0.894 | 0.085 | 1,200 | 0.892 | 0.099 | 1,060 | 0.783 | 0.080 |

**Supplementary Figure 1.** Correlation between pupal body size (cm2) and weight. Linear models are fitted for the control, large response, and selected lines independently. Each data point represents an average for each imaged batch of pupae, number of pupae per batch are presented by circle size.

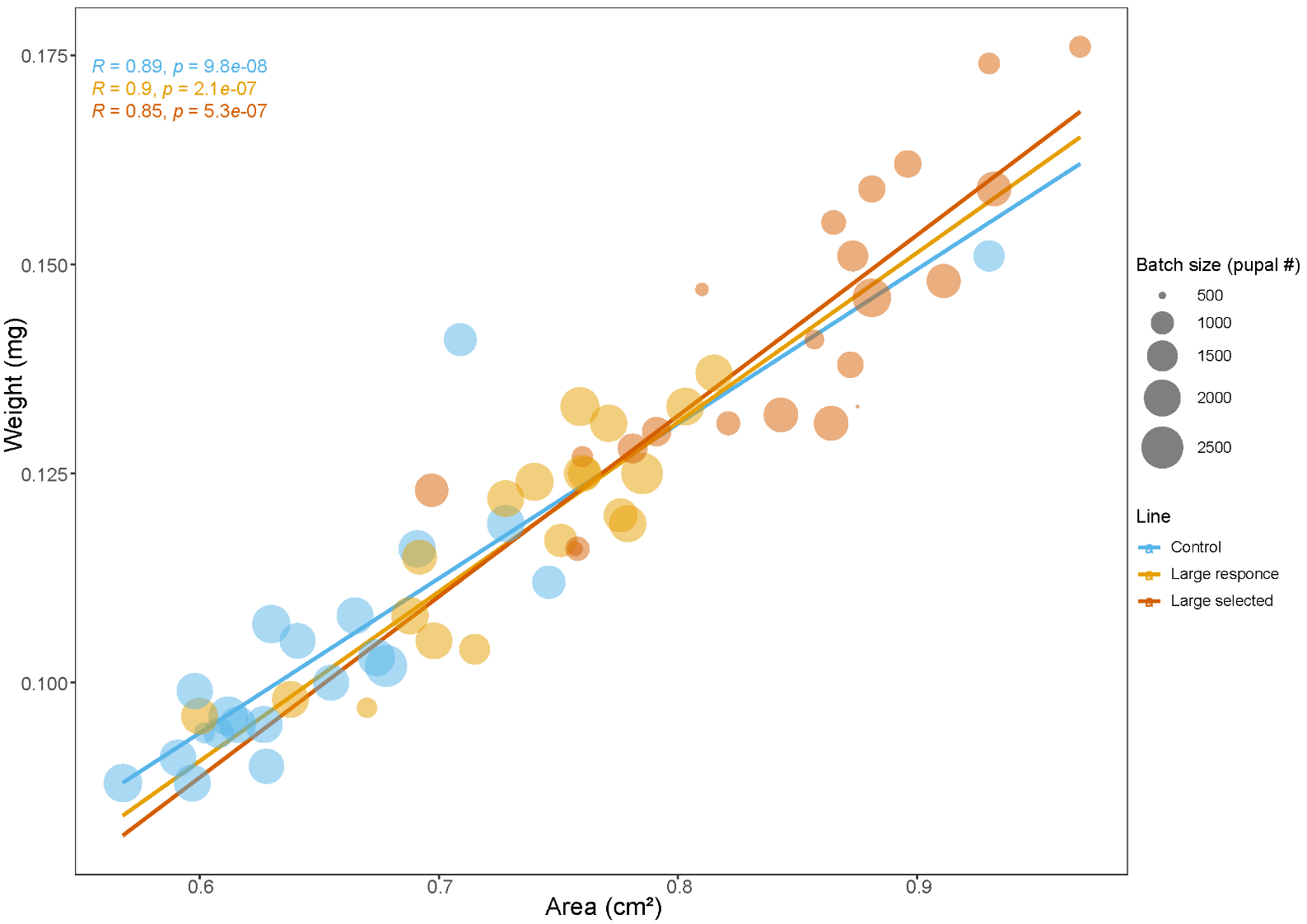

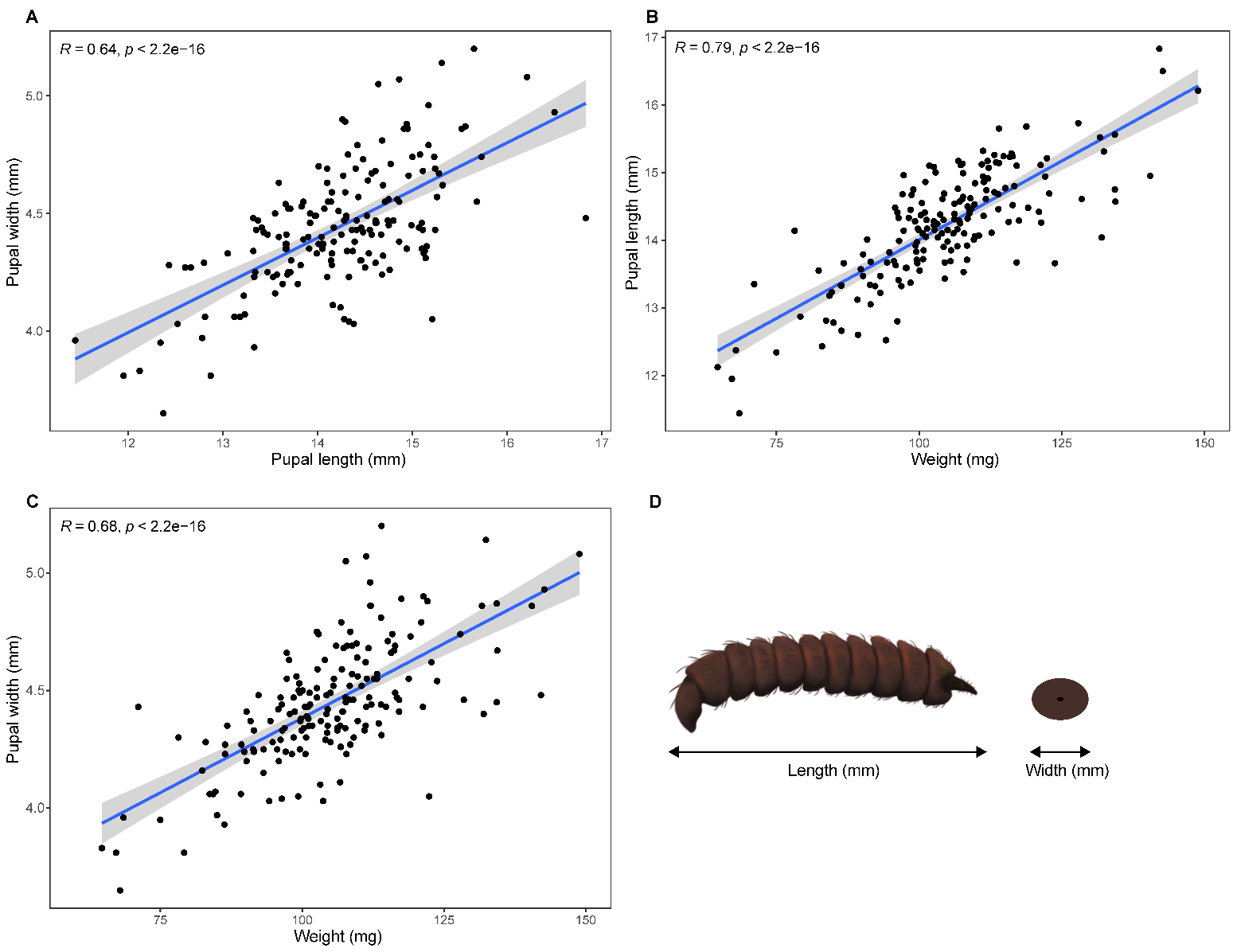

**Supplementary figure 2.** Correlations between pupal width - length (A), weight – length (B) and weight – width (C) showing allometric relationships between size metrics required for successful selection. Pupal width and length were recorded as shown in cross-sections in panel D.

**Supplementary figure 3.** Pupal body size phenotypes over seven generations of controlled rearing of a control and selected line for large pupal body size across three independent replicates. Mean phenotype of the starting base population is included alongside the control and selected lines.

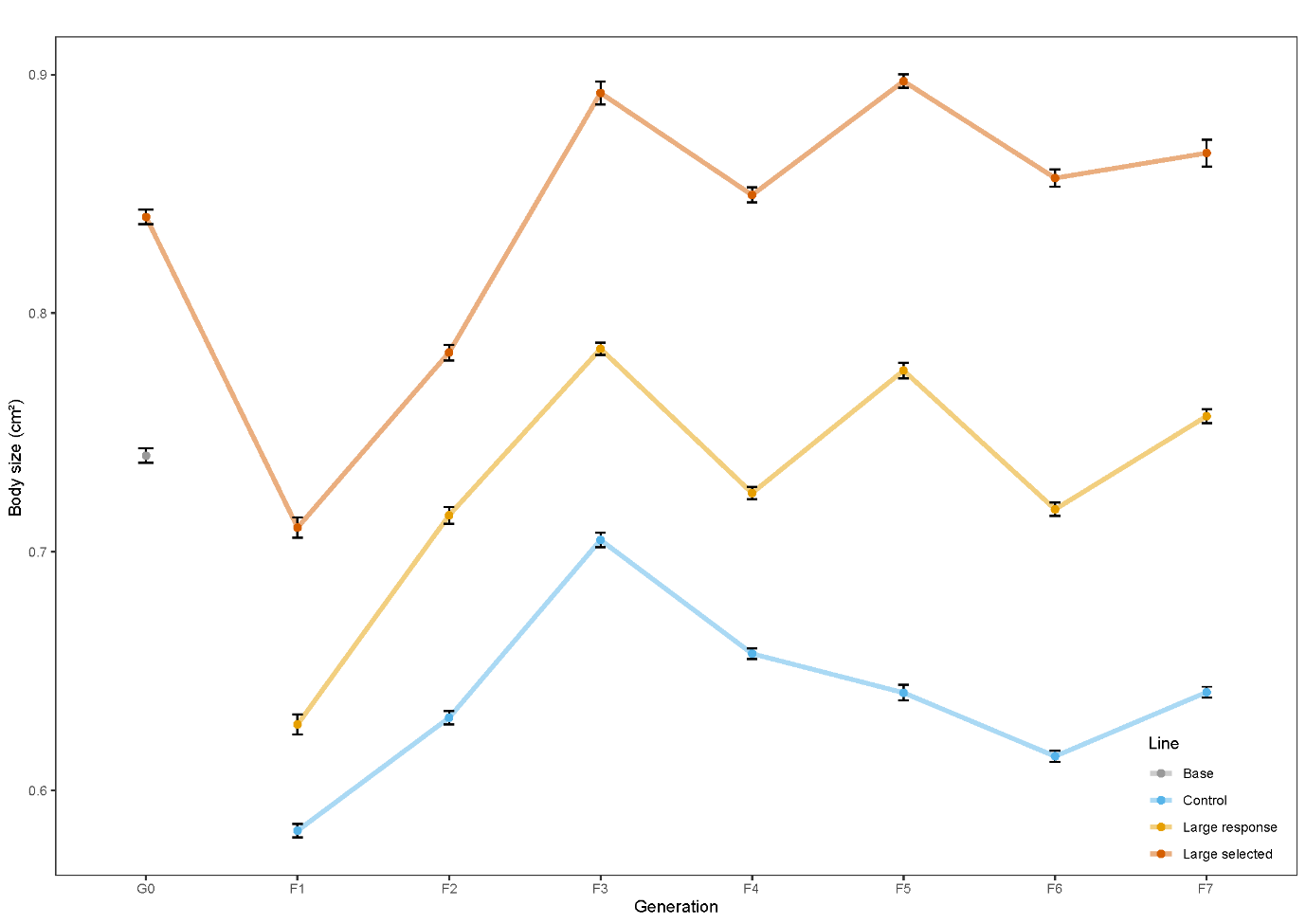

| **Supplementary Table 3.** Descriptive statistics for life-history traits of control and experimentally evolved lines at generation seven. Selection for increased pupal body size was performed in experimental lines over three independent replicates. Each line was maintained with an internal control (unselected). Average values for life-history traits and the standard deviation are presented for each replicate across various growth, development, nutrition, and breeding traits. Total number of sexed adults (female/total) is provided in addition to female proportion. | | | | | | |
| --- | --- | --- | --- | --- | --- | --- |
| **Life history trait** | **Replicate 01** | | **Replicate 02** | | **Replicate 03** | |
|  | **Control** | **Large** | **Control** | **Large** | **Control** | **Large** |
| **Growth traits** | | | | | | |
| Larval growth rate (mg/day) | 8.6 ± 5 | 11.1 ± 5.6 | 7.7 ± 3.3 | 9.2 ± 3.6 | 8.1 ± 4.8 | 8.8 ± 5.4 |
| Larval weight (mg) | 169.2 ± 26.7 | 202.8 ± 41.6 | 132.3 ± 25.9 | 143.0 ± 49.6 | 163.2 ± 29.5 | 198.1 ± 39.0 |
| Prepupal weight (mg) | 142.8 ± 24.7 | 177.8 ± 31.2 | 106.4 ± 21.1 | 129.0 ± 25.8 | 121.6 ± 19.9 | 147.0 ± 20.3 |
| Pupal weight (mg) | 95.5 ± 13.6 | 132.0 ± 26.3 | 86.3 ± 13.1 | 119.2 ± 15.4 | 98.7 ± 14.5 | 110.1 ± 11.5 |
| Female only | 105.4 ± 12.1 | 150.1 ± 23.2 | 87.7 ± 9 | 131.5 ± 14.4 | 108.5 ± 13.5 | 113.6 ± 14.2 |
| Male only | 93.3 ± 13.4 | 118.8 ± 19.3 | 87.6 ± 13.6 | 131.5 ± 14.1 | 94.4 ± 11.6 | 109.8 ± 10.1 |
| **Development rates** | | | | | | |
| Larval development rate (day-13 larval %) | 98.1 ± 0.4 | 84.8 ± 4.1 | 97.5 ± 1.4 | 94.2 ± 1.5 | 93.9 ± 2.3 | 94.9 ± 4.5 |
| Pupal development rate (hours) | 175 ± 21.7 | 178.3 ± 30.9 | 169.2 ± 13.7 | 175.2 ± 16.3 | 172.3 ± 14.2 | 169.2 ± 18.3 |
| Female only | 179.7 ± 13.9 | 189.6 ± 26.4 | 174.2 ± 11.6 | 176.7 ± 15.2 | 173.2 ± 14.5 | 174.3 ± 16.5 |
| Male only | 173.9 ± 23 | 169.8 ± 31.4 | 168.5 ± 13.9 | 174.9 ± 16.7 | 171.8 ± 14.2 | 168.5 ± 18.5 |
| **Nutrition traits** | | | | | | |
| Protein content (mg/L) | 18.6 ± 1.8 | 24.1 ± 1.7 | 13.9 ± 1.5 | 17.8 ± 2.8 | 21.3 ± 5.7 | 20.8 ± 6.2 |
| **Breeding traits** | | | | | | |
| Female proportion (%) | 21.5 (28/130) | 43 (64/149) | 12.3 (7/57) | 18.3 (13/71) | 36 (32/89) | 12.4 (10/81) |
| Ovary weight (mg) | 1.9 ± 0.8 | 4.1 ± 1.7 | 2.1 ± 1.2 | 2.8 ± 0.9 | 5.3 ± 2.5 | 5.32 ± 2.1 |

| **Supplementary table 4.** Linear Mixed-effect Model statistical analysis of sexed-pupal weights. Fixed effects include replicate, line, day, sex and interaction terms. Random effect of tray nested within each replicate is also characterised. | | | |
| --- | --- | --- | --- |
| **Fixed effect** | ***F*** | ***df*** | ***p*-value** |
| Line | 87.5 | 1 | 1.70E-06 |
| Sex | 129.7 | 1 | < 2.2e-16 |
| Line*Sex | 10.4 | 1 | 0.00133 |
| **Random effect** | **Variance** | **Variance (%)** | **Std. Dev** |
| Replicate | 0.005 | 0.84 | 0.229 |
| Tray:Replicate | 0.053 | 9.09 | 0.07 |
| Residuals | 0.521 | 90.07 | 0.722 |

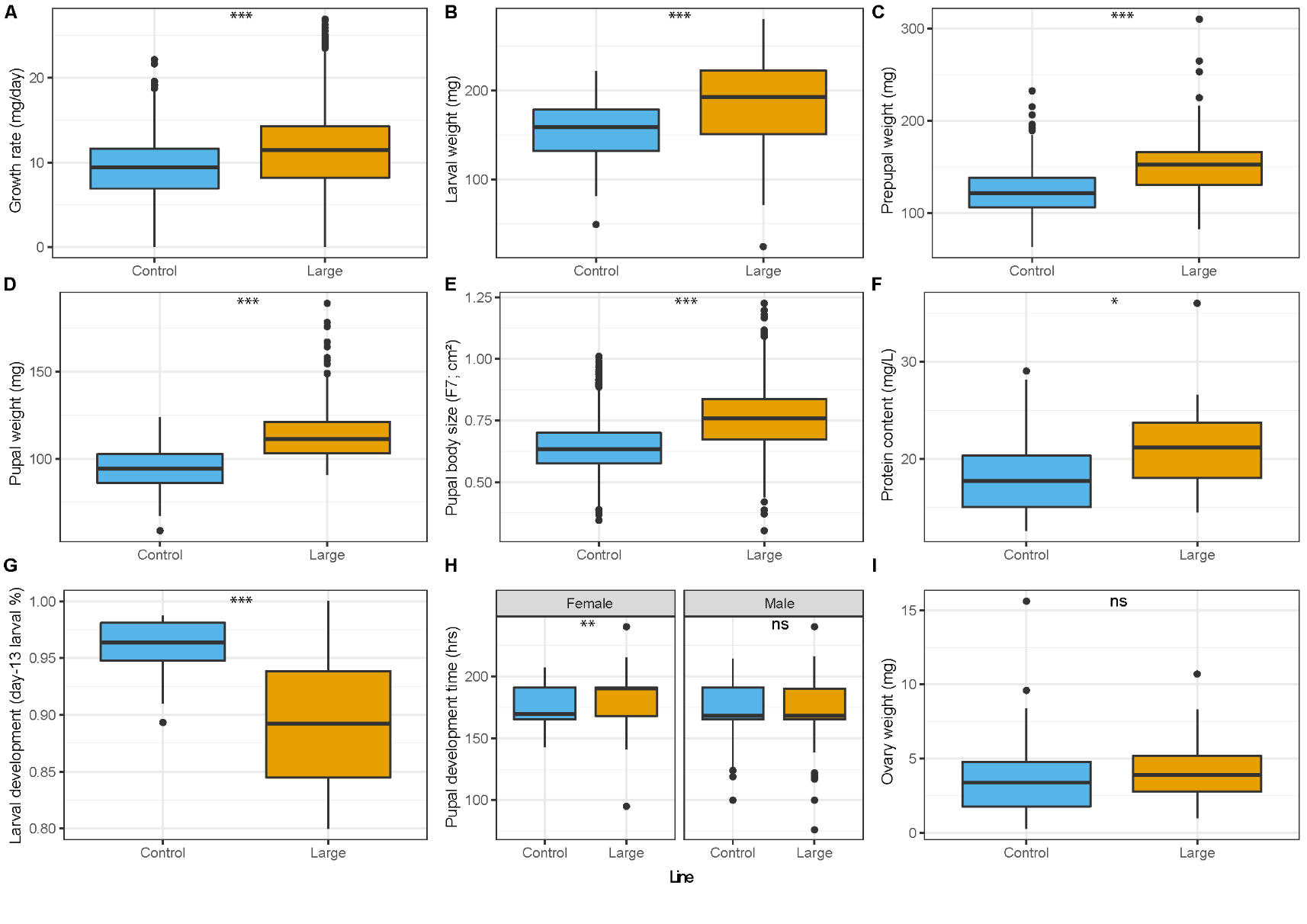

**Supplementary figure 4.** Pairwise comparisons between the control (blue) and large-selected (yellow) lines from the seventh generation of all three experiments. Measurements include Growth rate (mg/day) (A), day-12 larval weight (mg) (B), day-21 prepupal weight (mg) (C), day-18 pupal weight (mg) (D), pupal body size (cm2) (E), larval protein content (mg/larvae) (F), larval development (%) (G), pupal development (hours) (H) and adult ovary weight (mg) (I). Results comparing lines across replicates include t-test and Wilcoxon tests for traits shown (ns > 0.05, ,* < 0.05, ** < 0.01, *** < 0.001).

| **Supplementary table** **5**. Daily larval growth curve data for both control and selected lines at generation seven. Data for replicates are pooled into averages and standard deviation is provided. | | | | |
| --- | --- | --- | --- | --- |
| **Treatment** | **Day** | ***N*** | **Weight (mg)** | **SD** |
| Control | 5 | 200 | 4.989 | 6.044 |
| Large | 5 | 200 | 10.417 | 11.13 |
| Control | 6 | 200 | 12.009 | 7.49 |
| Large | 6 | 200 | 23.938 | 26.51 |
| Control | 7 | 225 | 29.45 | 15.89 |
| Large | 7 | 250 | 54.781 | 50.77 |
| Control | 8 | 250 | 53.571 | 17.87 |
| Large | 8 | 250 | 68.31 | 39.33 |
| Control | 9 | 250 | 109.228 | 29.35 |
| Large | 9 | 250 | 141.105 | 45.73 |
| Control | 10 | 250 | 96.429 | 28.68 |
| Large | 10 | 250 | 133.759 | 43.23 |
| Control | 11 | 250 | 110.088 | 20.84 |
| Large | 11 | 250 | 128.794 | 39.03 |
| Control | 12 | 250 | 155.726 | 31.51 |
| Large | 12 | 250 | 182.974 | 50.4 |
| Control | 13 | 250 | 141.83 | 30.17 |
| Large | 13 | 250 | 172.701 | 37.85 |
| Control | 14 | 250 | 132.629 | 25.99 |
| Large | 14 | 250 | 160.563 | 34.73 |
| Control | 15 | 250 | 185.456 | 40.02 |
| Large | 15 | 250 | 209.792 | 48.08 |
| Control | 16 | 250 | 174.576 | 36.86 |
| Large | 16 | 250 | 199.929 | 44.22 |
| Control | 17 | 250 | 154.482 | 31.64 |
| Large | 17 | 248 | 178.153 | 33.58 |
| Control | 18 | 250 | 169.076 | 39.72 |
| Large | 18 | 223 | 201.427 | 49.37 |
| Control | 19 | 239 | 188.586 | 52.14 |
| Large | 19 | 227 | 216.785 | 50.76 |
| Control | 20 | 221 | 187.604 | 56.53 |
| Large | 20 | 168 | 214.178 | 53.59 |

| **Supplementary table 6.** Linear Mixed-effect Model statistical analysis of daily measured larval weights. Fixed effects of line and day including the interaction between lines and days. Random effects of replicate and rearing trays nested within each replicate is also recorded. | | | |
| --- | --- | --- | --- |
| **Fixed effect** | ***F*** | ***df*** | ***p*-value** |
| Line | 46.11 | 9 | 8.00E-05 |
| Day | 1594.91 | 7558 | < 2.2e-16 |
| Line*Day | 6.09 | 7559 | 6.92E-13 |
| **Random effect** | **Variance** | **Variance (%)** | **Std. Dev** |
| Replicate | 269.92 | 17.44 | 16.429 |
| Tray:Replicate | 49.23 | 3.18 | 7.02 |
| Residuals | 1228.51 | 79.38 | 35.05 |

| **Supplementary table 7.** Linear Mixed-effect Model statistical analysis of protein proportions. Fixed effects of replicate and line and interactions between replicates and lines. Random effect of tray nested within each experiment is also characterised. | | | |
| --- | --- | --- | --- |
| **Fixed effect** | ***F*** | ***df*** | ***p*-value** |
| Line | 15.26 | 1 | 2.63E-04 |
| **Random effect** | **Variance** | **Variance (%)** | **Std. Dev** |
| Replicate | 0.005 | 41.45 | 0.05 |
| Tray:Replicate | 0.002 | 18.73 | 0.07 |
| Residuals | 0.005 | 39.82 | 0.07 |

| **Supplementary table 8.** Larval development rate of control and large-selected lines across all replicates at generation seven. Development assessed using average larval proportion sampled from rearing trays. Statistical analysis performed using a two-proportions-z-test at each day sampled between the two lines. | | | | |
| --- | --- | --- | --- | --- |
| **Day** | **Larval proportion (control; %)** | **Larval proportion (large; %)** | ***X^2^*** | ***p*-value** |
| 5 | 100.0 | 100.0 | NA | NA |
| 6 | 100.0 | 100.0 | NA | NA |
| 7 | 100.0 | 100.0 | NA | NA |
| 8 | 100.0 | 100.0 | NA | NA |
| 9 | 100.0 | 100.0 | NA | NA |
| 10 | 100.0 | 100.0 | NA | NA |
| 11 | 100.0 | 100.0 | NA | NA |
| 12 | 99.6 | 96.7 | 5.02 | 0.0250 |
| 13 | 95.9 | 89.1 | 5.39 | 0.0203 |
| 14 | 93.6 | 81.0 | 36.52 | 1.51E-09 |
| 15 | 80.1 | 72.5 | 5.13 | 0.0235 |
| 16 | 57.5 | 60.6 | 0.32 | 0.5725 |
| 17 | 43.3 | 48.8 | 0.59 | 0.4439 |
| 18 | 21.3 | 25.7 | 0.03 | 0.8667 |
| 19 | 7.1 | 18.8 | 37.10 | 1.12E-09 |
| 20 | 7.3 | 12.9 | 16.79 | 4.18E-05 |
| 21 | 3.9 | 7.6 | 8.96E-32 | 1 |
| 22 | 1.6 | 2.9 | 0.73 | 0.3926 |
| 23 | 1.4 | 2.0 | 0.00 | 0.9643 |
| 24 | 1.9 | 2.3 | 11.82 | 0.0006 |
| 25 | 0.8 | 0.9 | 0.07 | 0.7955 |

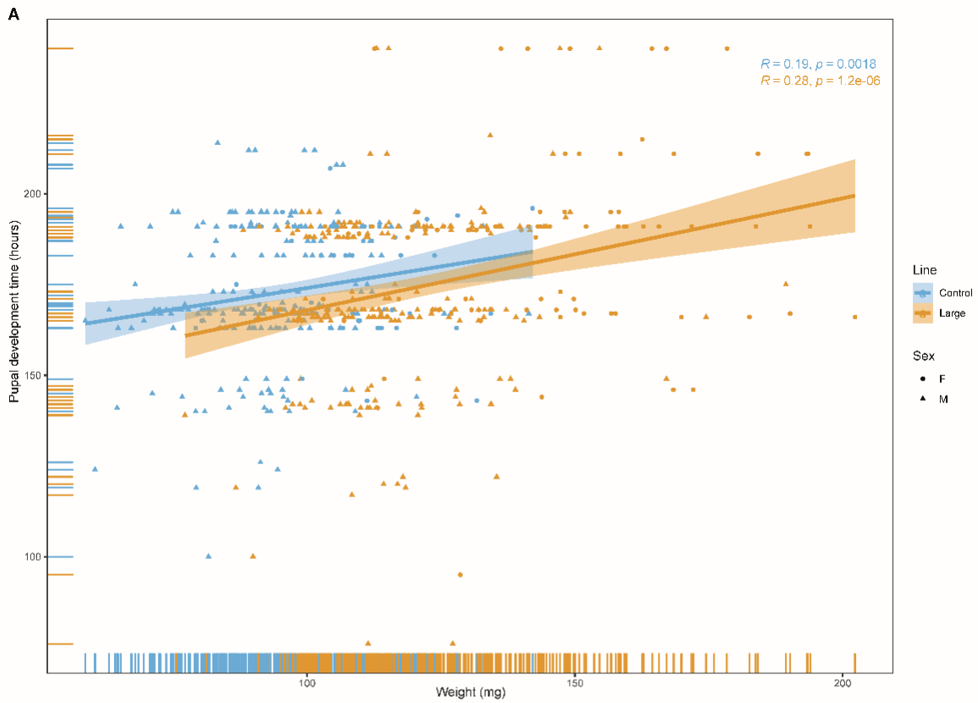

**Supplementary Figure 5** Pupal development time and weight relationships at generation seven. Correlation analysis is performed within the control and large-selected line showing a positive relationship, with increasing weight, pupal development is prolonged (A).

**Supplementary Table 9** Samples used for whole genome sequencing.

| **ID** | **Treatment** | **Generation** | **Sex** | **Replicate** |
| --- | --- | --- | --- | --- |
| CAM006241 | BASE | G0 | Male | R3 |
| CAM006242 | BASE | G0 | Female | R3 |
| CAM006244 | BASE | G0 | Female | R3 |
| CAM006245 | BASE | G0 | Male | R3 |
| CAM006246 | BASE | G0 | Female | R3 |
| CAM006248 | BASE | G0 | Female | R3 |
| CAM006259 | LARGE | G7 | Male | R3 |
| CAM006260 | LARGE | G7 | Female | R3 |
| CAM006261 | LARGE | G7 | Male | R3 |
| CAM006262 | LARGE | G7 | Female | R3 |
| CAM006263 | LARGE | G7 | Male | R3 |
| CAM006264 | LARGE | G7 | Female | R3 |
| CAM006265 | LARGE | G7 | Male | R3 |
| CAM006266 | LARGE | G7 | Female | R3 |
| CAM006268 | CONTROL | G7 | Female | R3 |
| CAM006269 | CONTROL | G7 | Male | R3 |
| CAM006278 | BASE | G0 | Female | R2 |
| CAM006280 | BASE | G0 | Female | R2 |
| CAM006281 | BASE | G0 | Male | R2 |
| CAM006283 | BASE | G0 | Male | R2 |
| CAM006287 | CONTROL | G7 | Male | R2 |
| CAM006289 | CONTROL | G7 | Male | R2 |
| CAM006290 | CONTROL | G7 | Female | R2 |
| CAM006298 | LARGE | G7 | Female | R2 |
| CAM006305 | BASE | G0 | Male | R1 |
| CAM006307 | BASE | G0 | Male | R1 |
| CAM006308 | BASE | G0 | Female | R1 |
| CAM006309 | BASE | G0 | Male | R1 |
| CAM006313 | CONTROL | G7 | Male | R1 |
| CAM006319 | CONTROL | G7 | Male | R1 |
| CAM006322 | LARGE | G7 | Female | R1 |
| CAM006323 | LARGE | G7 | Male | R1 |
| CAM006327 | LARGE | G7 | Male | R1 |
| CAM006328 | LARGE | G7 | Female | R1 |
| CAM006329 | LARGE | G7 | Male | R1 |
| CAM006701 | BASE | G0 | Male | R1 |
| CAM006702 | BASE | G0 | Female | R1 |
| CAM006703 | BASE | G0 | Male | R1 |
| CAM006704 | BASE | G0 | Female | R1 |
| CAM006706 | BASE | G0 | Female | R1 |
| CAM006707 | BASE | G0 | Male | R1 |
| CAM006708 | BASE | G0 | Female | R1 |
| CAM006711 | BASE | G0 | Male | R1 |
| CAM006712 | BASE | G0 | Female | R1 |
| CAM006713 | BASE | G0 | Male | R1 |
| CAM006714 | BASE | G0 | Female | R1 |
| CAM006716 | BASE | G0 | Female | R1 |
| CAM006717 | BASE | G0 | Male | R1 |
| CAM006718 | BASE | G0 | Female | R1 |
| CAM006719 | BASE | G0 | Male | R1 |
| CAM006720 | BASE | G0 | Female | R1 |
| CAM006721 | BASE | G0 | Male | R1 |
| CAM006722 | BASE | G0 | Female | R1 |
| CAM006724 | BASE | G0 | Female | R1 |
| CAM006725 | BASE | G0 | Male | R1 |
| CAM006726 | BASE | G0 | Female | R1 |
| CAM006727 | BASE | G0 | Male | R1 |
| CAM006728 | BASE | G0 | Female | R1 |
| CAM006729 | BASE | G0 | Male | R1 |
| CAM006730 | BASE | G0 | Female | R1 |
| CAM006731 | BASE | G0 | Male | R1 |
| CAM006732 | BASE | G0 | Female | R1 |
| CAM006741 | BASE | G0 | Male | R2 |
| CAM006742 | BASE | G0 | Female | R2 |
| CAM006743 | BASE | G0 | Male | R2 |
| CAM006744 | BASE | G0 | Female | R2 |
| CAM006745 | BASE | G0 | Male | R2 |
| CAM006746 | BASE | G0 | Female | R2 |
| CAM006747 | BASE | G0 | Male | R2 |
| CAM006748 | BASE | G0 | Female | R2 |
| CAM006757 | BASE | G0 | Male | R2 |
| CAM006761 | BASE | G0 | Male | R2 |
| CAM006765 | BASE | G0 | Male | R3 |
| CAM006766 | BASE | G0 | Female | R3 |
| CAM006767 | BASE | G0 | Male | R3 |
| CAM006768 | BASE | G0 | Female | R3 |
| CAM006769 | BASE | G0 | Male | R3 |
| CAM006770 | BASE | G0 | Female | R3 |
| CAM006771 | BASE | G0 | Male | R3 |
| CAM006772 | BASE | G0 | Female | R3 |
| CAM006773 | BASE | G0 | Male | R3 |
| CAM006774 | BASE | G0 | Female | R3 |
| CAM006776 | BASE | G0 | Female | R3 |
| CAM006777 | BASE | G0 | Male | R3 |
| CAM006782 | BASE | G0 | Female | R3 |
| CAM006788 | BASE | G0 | Female | R3 |
| CAM006793 | BASE | G0 | Male | R3 |
| CAM006796 | BASE | G0 | Female | R3 |
| CAM006799 | CONTROL | G7 | Male | R1 |
| CAM006803 | CONTROL | G7 | Male | R1 |
| CAM006805 | CONTROL | G7 | Male | R1 |
| CAM006807 | CONTROL | G7 | Male | R1 |
| CAM006809 | CONTROL | G7 | Male | R1 |
| CAM006811 | CONTROL | G7 | Male | R1 |
| CAM006812 | CONTROL | G7 | Female | R1 |
| CAM006813 | CONTROL | G7 | Male | R1 |
| CAM006814 | CONTROL | G7 | Female | R1 |
| CAM006815 | CONTROL | G7 | Male | R1 |
| CAM006816 | CONTROL | G7 | Female | R1 |
| CAM006817 | CONTROL | G7 | Male | R1 |
| CAM006819 | CONTROL | G7 | Male | R1 |
| CAM006820 | CONTROL | G7 | Female | R1 |
| CAM006825 | CONTROL | G7 | Male | R1 |
| CAM006829 | CONTROL | G7 | Male | R2 |
| CAM006830 | CONTROL | G7 | Female | R2 |
| CAM006831 | CONTROL | G7 | Male | R2 |
| CAM006832 | CONTROL | G7 | Female | R2 |
| CAM006833 | CONTROL | G7 | Male | R2 |
| CAM006834 | CONTROL | G7 | Female | R2 |
| CAM006835 | CONTROL | G7 | Male | R2 |
| CAM006836 | CONTROL | G7 | Female | R2 |
| CAM006841 | CONTROL | G7 | Male | R2 |
| CAM006842 | CONTROL | G7 | Female | R2 |
| CAM006843 | CONTROL | G7 | Male | R2 |
| CAM006844 | CONTROL | G7 | Female | R2 |
| CAM006847 | CONTROL | G7 | Male | R2 |
| CAM006848 | CONTROL | G7 | Female | R2 |
| CAM006850 | CONTROL | G7 | Female | R2 |
| CAM006853 | CONTROL | G7 | Male | R2 |
| CAM006854 | CONTROL | G7 | Female | R2 |
| CAM006855 | CONTROL | G7 | Male | R2 |
| CAM006856 | CONTROL | G7 | Female | R2 |
| CAM006857 | CONTROL | G7 | Male | R2 |
| CAM006858 | CONTROL | G7 | Female | R2 |
| CAM006859 | CONTROL | G7 | Male | R2 |
| CAM006860 | CONTROL | G7 | Female | R2 |
| CAM006862 | CONTROL | G7 | Female | R3 |
| CAM006864 | CONTROL | G7 | Female | R3 |
| CAM006867 | CONTROL | G7 | Male | R3 |
| CAM006877 | CONTROL | G7 | Male | R3 |
| CAM006884 | CONTROL | G7 | Female | R3 |
| CAM006885 | CONTROL | G7 | Male | R3 |
| CAM006890 | CONTROL | G7 | Female | R3 |
| CAM006894 | LARGE | G7 | Female | R1 |
| CAM006895 | LARGE | G7 | Male | R1 |
| CAM006896 | LARGE | G7 | Female | R1 |
| CAM006897 | LARGE | G7 | Male | R1 |
| CAM006898 | LARGE | G7 | Female | R1 |
| CAM006899 | LARGE | G7 | Male | R1 |
| CAM006900 | LARGE | G7 | Female | R1 |
| CAM006901 | LARGE | G7 | Male | R1 |
| CAM006902 | LARGE | G7 | Female | R1 |
| CAM006903 | LARGE | G7 | Male | R1 |
| CAM006904 | LARGE | G7 | Female | R1 |
| CAM006905 | LARGE | G7 | Male | R1 |
| CAM006906 | LARGE | G7 | Female | R1 |
| CAM006907 | LARGE | G7 | Male | R1 |
| CAM006908 | LARGE | G7 | Female | R1 |
| CAM006909 | LARGE | G7 | Male | R1 |
| CAM006911 | LARGE | G7 | Male | R1 |
| CAM006912 | LARGE | G7 | Female | R1 |
| CAM006913 | LARGE | G7 | Male | R1 |
| CAM006914 | LARGE | G7 | Female | R1 |
| CAM006917 | LARGE | G7 | Male | R1 |
| CAM006918 | LARGE | G7 | Female | R1 |
| CAM006919 | LARGE | G7 | Male | R1 |
| CAM006920 | LARGE | G7 | Female | R1 |
| CAM006921 | LARGE | G7 | Male | R1 |
| CAM006922 | LARGE | G7 | Female | R1 |
| CAM006924 | LARGE | G7 | Female | R1 |
| CAM006926 | LARGE | G7 | Female | R2 |
| CAM006930 | LARGE | G7 | Female | R2 |
| CAM006931 | LARGE | G7 | Male | R2 |
| CAM006932 | LARGE | G7 | Female | R2 |
| CAM006936 | LARGE | G7 | Female | R2 |
| CAM006940 | LARGE | G7 | Female | R2 |
| CAM006941 | LARGE | G7 | Male | R2 |
| CAM006942 | LARGE | G7 | Female | R2 |
| CAM006944 | LARGE | G7 | Female | R2 |
| CAM006945 | LARGE | G7 | Male | R2 |
| CAM006946 | LARGE | G7 | Female | R2 |
| CAM006948 | LARGE | G7 | Female | R2 |
| CAM006950 | LARGE | G7 | Female | R2 |
| CAM006951 | LARGE | G7 | Male | R2 |
| CAM006952 | LARGE | G7 | Female | R2 |
| CAM006954 | LARGE | G7 | Female | R2 |
| CAM006958 | LARGE | G7 | Female | R3 |
| CAM006965 | LARGE | G7 | Male | R3 |
| CAM006966 | LARGE | G7 | Female | R3 |
| CAM006967 | LARGE | G7 | Male | R3 |
| CAM006968 | LARGE | G7 | Female | R3 |
| CAM006969 | LARGE | G7 | Male | R3 |
| CAM006970 | LARGE | G7 | Female | R3 |
| CAM006971 | LARGE | G7 | Male | R3 |
| CAM006972 | LARGE | G7 | Female | R3 |
| CAM006973 | LARGE | G7 | Male | R3 |
| CAM006977 | LARGE | G7 | Male | R3 |
| CAM006978 | LARGE | G7 | Female | R3 |
| CAM006979 | LARGE | G7 | Male | R3 |
| CAM006982 | LARGE | G7 | Female | R3 |
| CAM006984 | LARGE | G7 | Female | R3 |
| CAM006985 | LARGE | G7 | Male | R3 |
| CAM006986 | LARGE | G7 | Female | R3 |
| CAM006987 | LARGE | G7 | Male | R3 |
| CAM006989 | CONTROL | G7 | Male | R1 |
| CAM006990 | CONTROL | G7 | Female | R1 |
| CAM006991 | CONTROL | G7 | Male | R1 |
| CAM006992 | CONTROL | G7 | Female | R1 |
| CAM006993 | CONTROL | G7 | Male | R1 |
| CAM006994 | CONTROL | G7 | Female | R1 |
| CAM006995 | BASE | G0 | Male | R2 |
| CAM006996 | CONTROL | G7 | Female | R1 |
| CAM006997 | BASE | G0 | Male | R2 |
| CAM006998 | CONTROL | G7 | Female | R1 |
| CAM006999 | BASE | G0 | Male | R2 |
| CAM007000 | CONTROL | G7 | Female | R1 |
| CAM007001 | BASE | G0 | Male | R2 |
| CAM007002 | CONTROL | G7 | Female | R1 |
| CAM007003 | BASE | G0 | Male | R2 |
| CAM007004 | CONTROL | G7 | Female | R1 |
| CAM007005 | BASE | G0 | Male | R2 |
| CAM007006 | CONTROL | G7 | Female | R1 |
| CAM007007 | BASE | G0 | Male | R2 |
| CAM007008 | CONTROL | G7 | Female | R1 |
| CAM007009 | BASE | G0 | Male | R2 |
| CAM007010 | CONTROL | G7 | Female | R1 |
| CAM007011 | BASE | G0 | Male | R3 |
| CAM007012 | CONTROL | G7 | Female | R1 |
| CAM007013 | BASE | G0 | Male | R3 |
| CAM007014 | BASE | G0 | Female | R2 |
| CAM007015 | BASE | G0 | Male | R3 |
| CAM007016 | BASE | G0 | Female | R2 |
| CAM007017 | BASE | G0 | Male | R3 |
| CAM007018 | BASE | G0 | Female | R2 |
| CAM007019 | CONTROL | G7 | Male | R2 |
| CAM007021 | CONTROL | G7 | Male | R2 |
| CAM007023 | CONTROL | G7 | Male | R2 |
| CAM007024 | BASE | G0 | Female | R2 |
| CAM007025 | LARGE | G7 | Male | R2 |
| CAM007026 | BASE | G0 | Female | R2 |
| CAM007027 | LARGE | G7 | Male | R2 |
| CAM007028 | BASE | G0 | Female | R2 |
| CAM007029 | LARGE | G7 | Male | R2 |
| CAM007030 | BASE | G0 | Female | R2 |
| CAM007031 | LARGE | G7 | Male | R2 |
| CAM007032 | BASE | G0 | Female | R2 |
| CAM007033 | LARGE | G7 | Male | R2 |
| CAM007034 | BASE | G0 | Female | R2 |
| CAM007035 | LARGE | G7 | Male | R2 |
| CAM007036 | BASE | G0 | Female | R2 |
| CAM007037 | LARGE | G7 | Male | R2 |
| CAM007038 | CONTROL | G7 | Female | R2 |
| CAM007039 | LARGE | G7 | Male | R2 |
| CAM007040 | CONTROL | G7 | Female | R2 |
| CAM007041 | LARGE | G7 | Male | R2 |
| CAM007042 | CONTROL | G7 | Female | R2 |
| CAM007043 | LARGE | G7 | Male | R2 |
| CAM007044 | LARGE | G7 | Female | R2 |
| CAM007045 | LARGE | G7 | Male | R2 |
| CAM007046 | LARGE | G7 | Female | R2 |
| CAM007047 | LARGE | G7 | Male | R2 |
| CAM007048 | LARGE | G7 | Female | R2 |
| CAM007049 | BASE | G0 | Male | R3 |
| CAM007050 | BASE | G0 | Female | R3 |
| CAM007051 | BASE | G0 | Male | R3 |
| CAM007052 | BASE | G0 | Female | R3 |
| CAM007053 | BASE | G0 | Male | R3 |
| CAM007054 | BASE | G0 | Female | R3 |
| CAM007055 | CONTROL | G7 | Male | R3 |
| CAM007056 | CONTROL | G7 | Female | R3 |
| CAM007057 | CONTROL | G7 | Male | R3 |
| CAM007058 | CONTROL | G7 | Female | R3 |
| CAM007059 | CONTROL | G7 | Male | R3 |
| CAM007060 | CONTROL | G7 | Female | R3 |
| CAM007061 | CONTROL | G7 | Male | R3 |
| CAM007062 | CONTROL | G7 | Female | R3 |
| CAM007063 | CONTROL | G7 | Male | R3 |
| CAM007064 | CONTROL | G7 | Female | R3 |
| CAM007065 | CONTROL | G7 | Male | R3 |
| CAM007066 | CONTROL | G7 | Female | R3 |
| CAM007067 | CONTROL | G7 | Male | R3 |
| CAM007068 | CONTROL | G7 | Female | R3 |
| CAM007069 | CONTROL | G7 | Male | R3 |
| CAM007070 | CONTROL | G7 | Female | R3 |
| CAM007071 | CONTROL | G7 | Male | R3 |
| CAM007072 | CONTROL | G7 | Female | R3 |
| CAM007073 | CONTROL | G7 | Male | R3 |
| CAM007074 | CONTROL | G7 | Female | R3 |
| CAM007075 | CONTROL | G7 | Male | R3 |
| CAM007076 | CONTROL | G7 | Female | R3 |
| CAM007077 | CONTROL | G7 | Male | R3 |
| CAM007078 | CONTROL | G7 | Female | R3 |
| CAM007079 | LARGE | G7 | Male | R3 |
| CAM007080 | LARGE | G7 | Female | R3 |
| CAM007081 | LARGE | G7 | Male | R3 |
| CAM007082 | LARGE | G7 | Female | R3 |
| CAM007083 | LARGE | G7 | Male | R3 |
| CAM007084 | LARGE | G7 | Female | R3 |

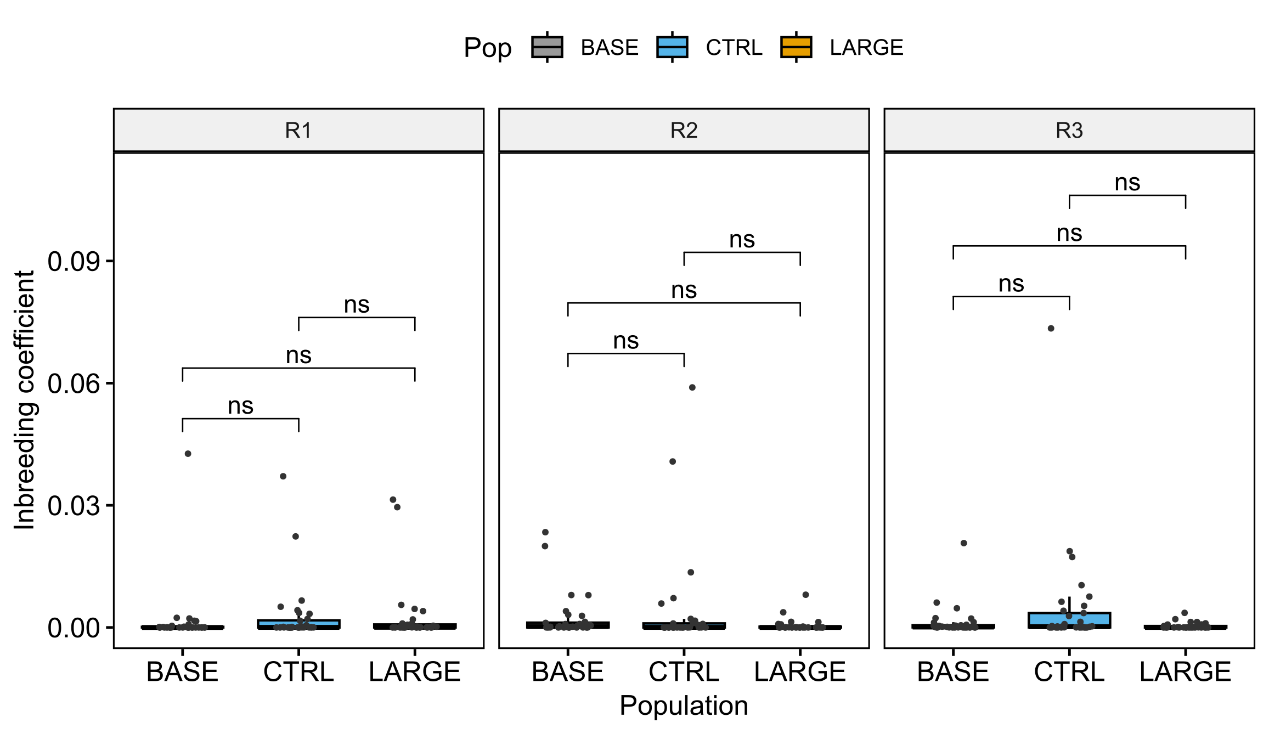

**Supplementary Figure 6** Inbreeding coefficient per individual of all lines in all replicates. Replicate 1, 2 and 3 are represented as “R1”, “R2” and “R3”, respectively.

**Supplementary Table 10** Average nucleotide diversity based on 20kb sliding windows for each population in each replicate.

| **Replicate** | **Line** | **Average pi** |
| --- | --- | --- |
| R1 | BASE | 0.0187 |
|  | CTRL | 0.0186 |
|  | LARGE | 0.0181 |
| R2 | BASE | 0.0188 |
|  | CTRL | 0.0180 |
|  | LARGE | 0.0158 |
| R3 | BASE | 0.0190 |
|  | CTRL | 0.0182 |
|  | LARGE | 0.0173 |

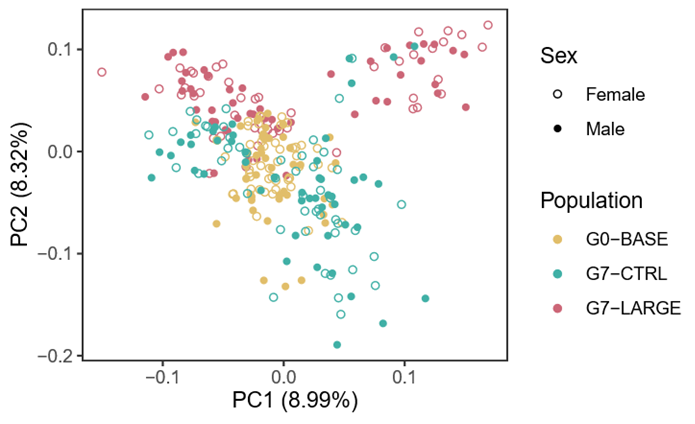
**A**

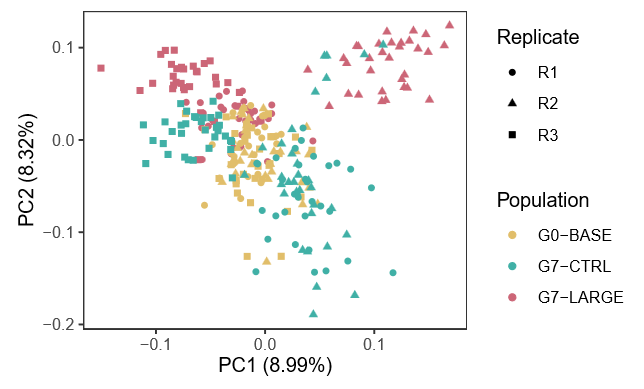

**B**

**Supplementary Figure 7** PCA plot divided by sex. Male and Female individuals do not cluster separately in any population.

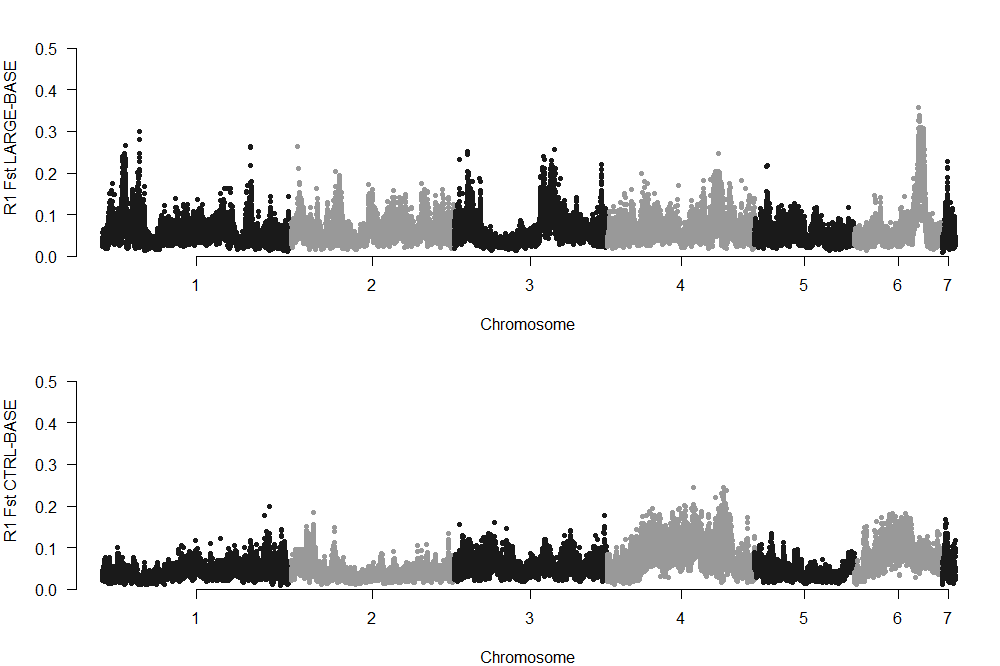

**B**

**A**

**C**

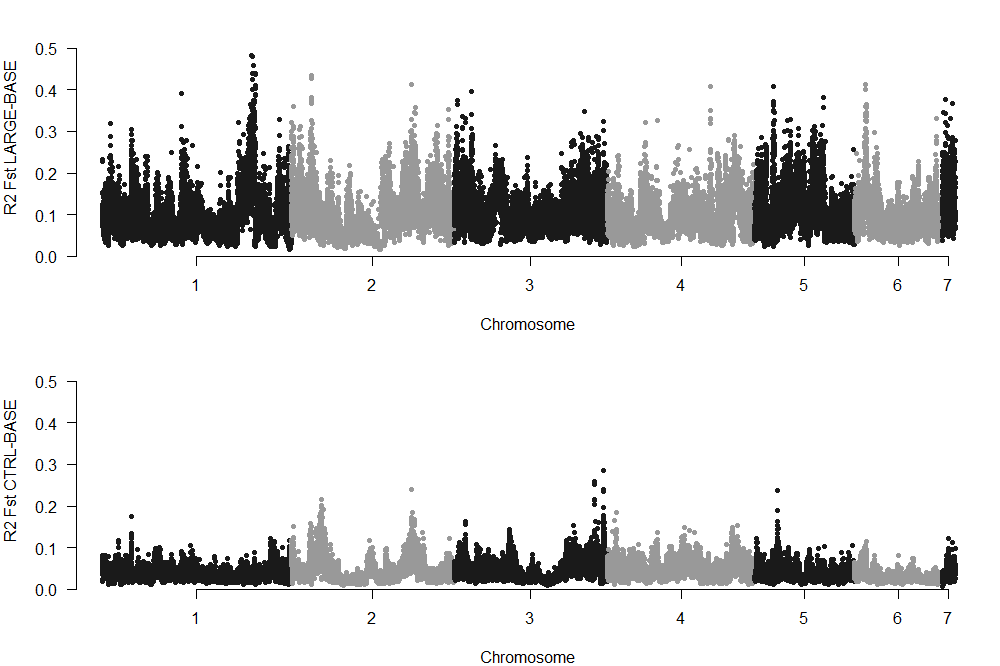

**D**

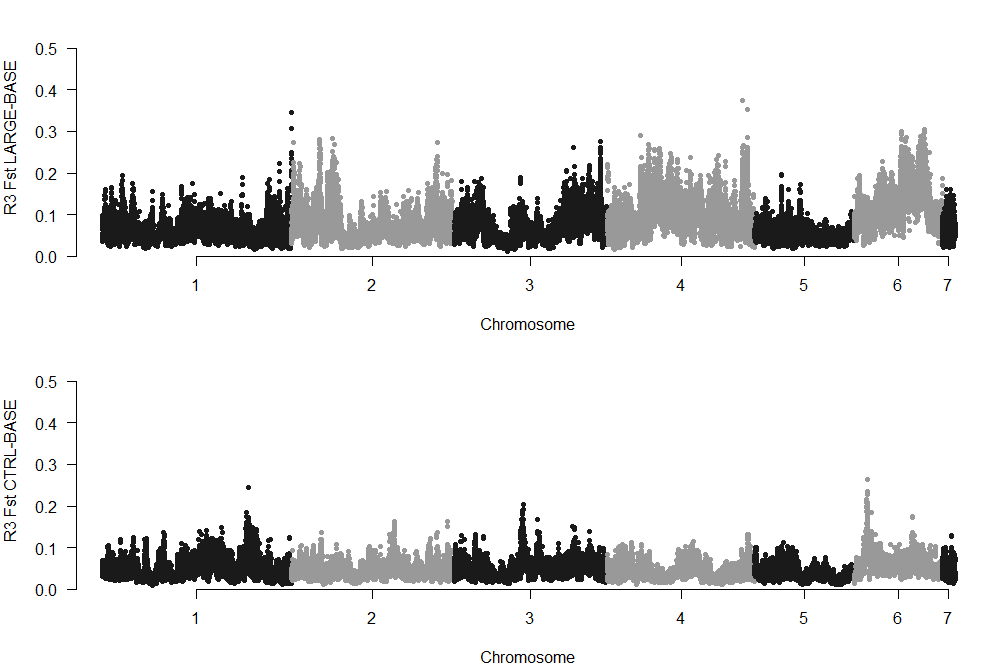

**F**

**E**

**Supplementary Figure 8** *F*_ST_ landscape of LARGE-BASE and CTRL-BASE in replicate 1 (**A**-**B**), 2 (**C**-**D**), and 3 (**E**-**F**).

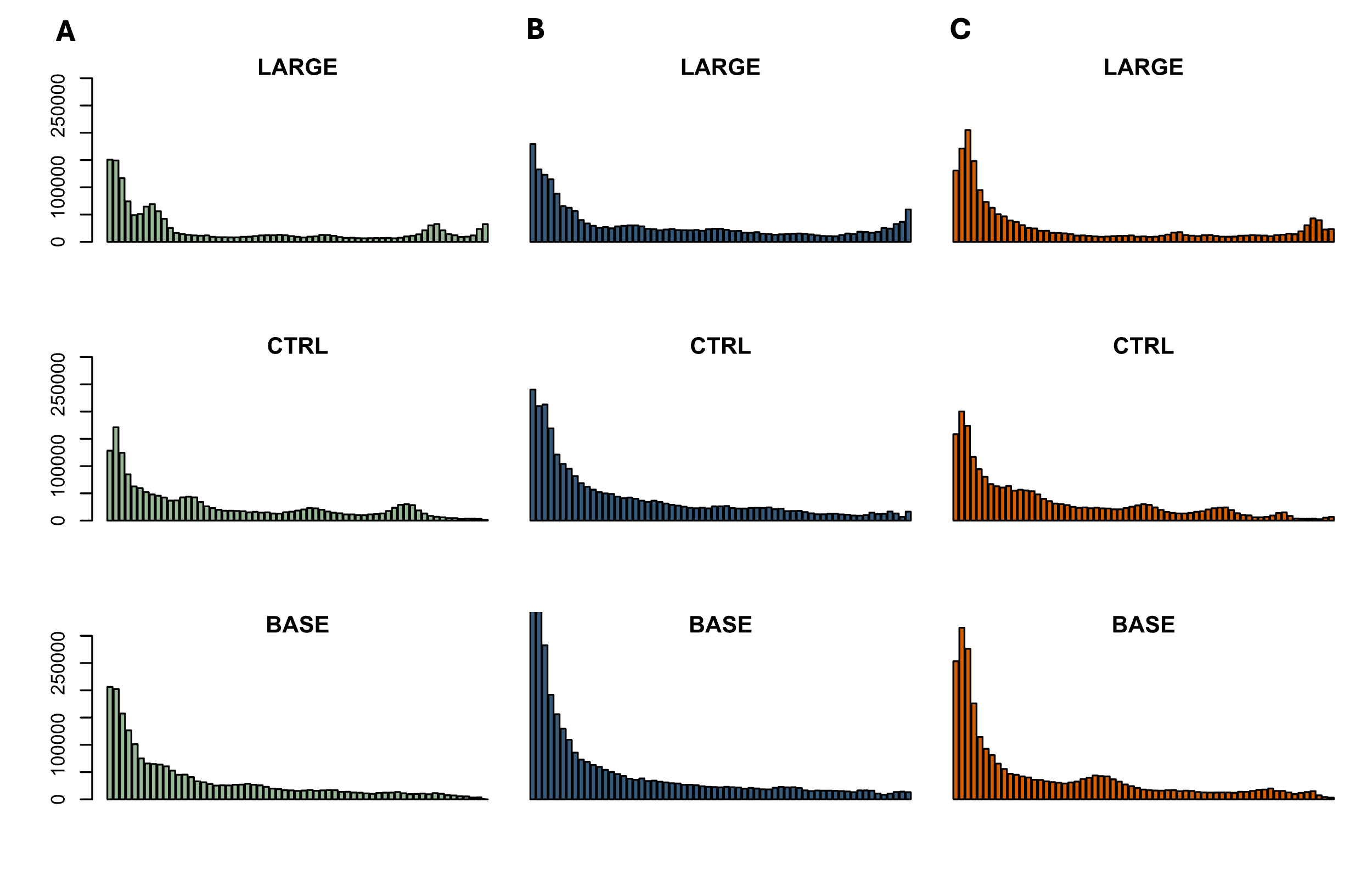

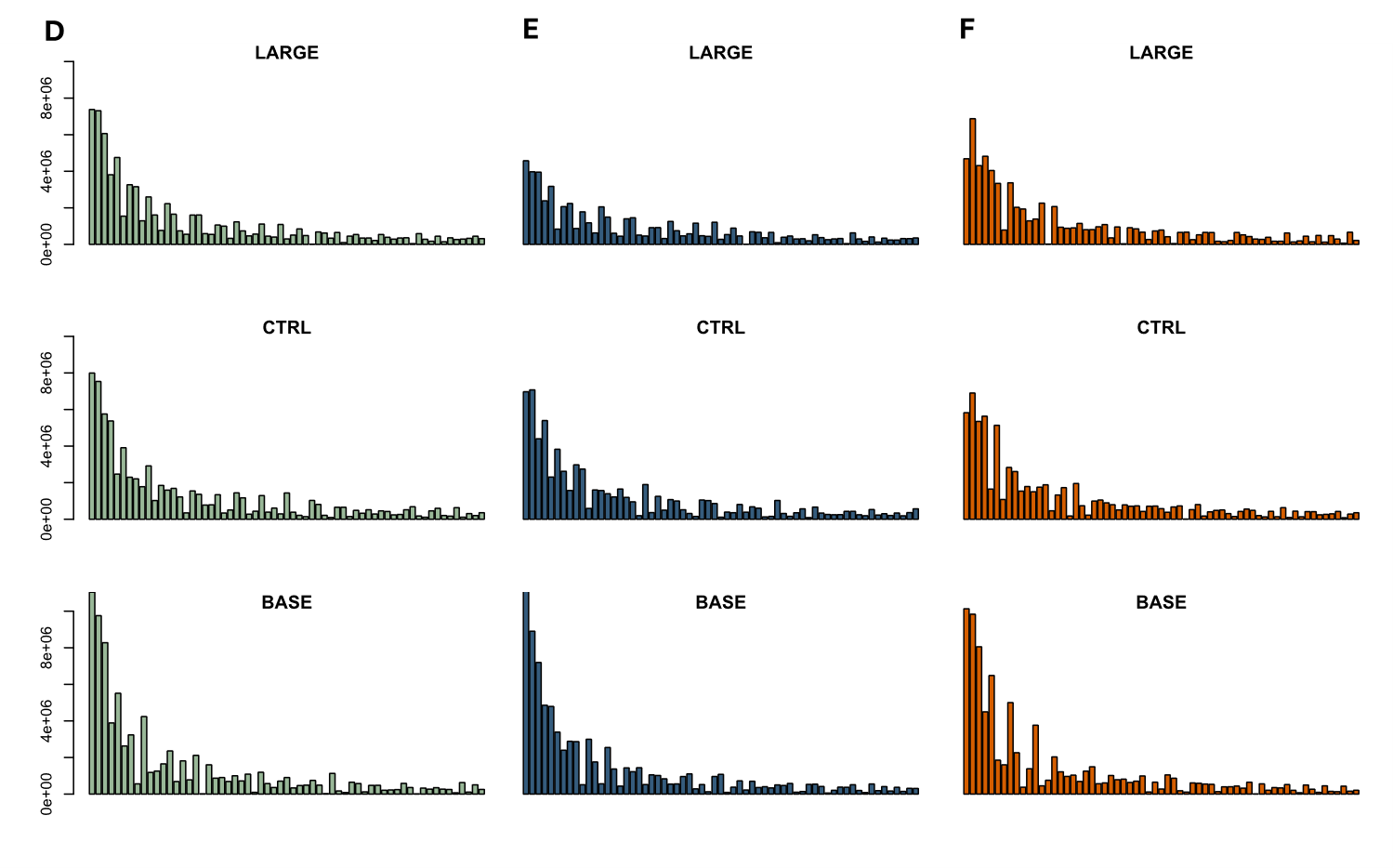

**Supplementary Figure 9** Shift in site frequency spectrum pattern of high divergence windows (**A**-**C**) and whole genome (**D**-**F**) in replicate 1-3, respectively. The X-axis represents derived allele frequency, and the Y-axis represents numbers of sites that fall within each category of alleles. The upper limit of Y-axis is set to be the same in each replicate for plotting convenience. An increase in high-frequency alleles between base and the selected population is seen in high divergence windows in all replicates, while only a decrease in low-frequency alleles is observed in the whole genome comparison.

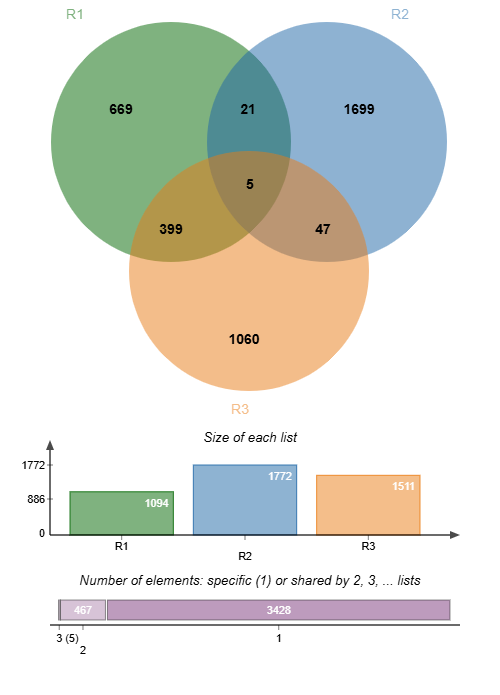

**Supplementary Figure 10** Venn diagram of shared sliding windows among all three replicates. Replicate 1 and 3 share a higher number of windows compared to each of them with replicate 2. Only 5 windows are shared in all three replicates and all of them locate in the same continuous peak region between 75160000-75840000bp on Chromosome 6.

**Supplementary Table 11** Candidate peak region and gene list

| **Chromosome** | **Region start** | **Region end** | **Gene start** | **Gene end** | **Gene ID** | **Gene name** |
| --- | --- | --- | --- | --- | --- | --- |

| S3 | 172720000 | 172780000 | 172760254 | 172766397 | LOC119651082 | *LOC119651082* |
| --- | --- | --- | --- | --- | --- | --- |
|  |  |  | 172772527 | 172775676 | LOC119651081 | *eIF4A-like* |
|  |  |  | 172726793 | 172756011 | LOC119650632 | *eIF4A* |
| S4 | 130560000 | 130700000 | 130680626 | 130680699 | Trnat-agu-6 | *Trnat-agu-6* |
|  |  |  | 130655462 | 130664874 | LOC119653877 | *CysRS* |
|  |  |  | 130686772 | 130700201 | LOC119654788 | *NiPp1* |
|  |  |  | 130665193 | 130680397 | LOC119653878 | *eIF3m* |
| S4 | 131380000 | 131460000 | 130933925 | 131470829 | LOC119653887 | *Pkc53E* |
| S6 | 75160000 | 75840000 | 75831949 | 75832987 | LOC119659542 | *LOC119659542* |
|  |  |  | 75817774 | 75831737 | LOC119659933 | *Nab2* |
|  |  |  | 75832020 | 75846504 | LOC119659541 | *Mps1* |
|  |  |  | 75619223 | 75812774 | LOC119660051 | *HiInR* |

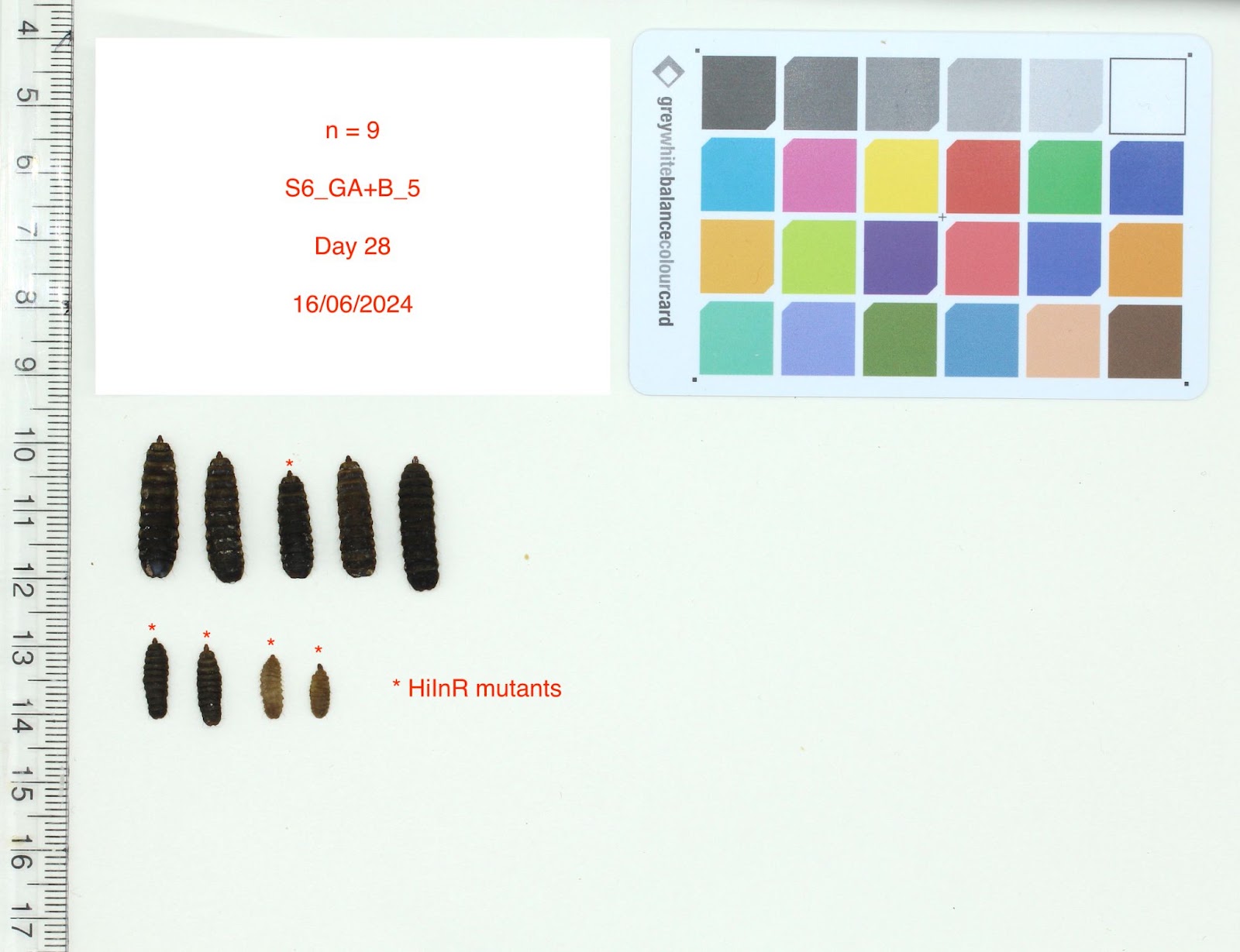

**Supplementary Figure 11** A *HiInR* CRISPR group with 5 mutants confirmed by genotyping. Exceptional small size and relatively earlier developmental stages (prepupal and larval stage) can be seen in the bottom four mutants. No phenotypic abnormality was observed in the top centre mutant, except smaller size.

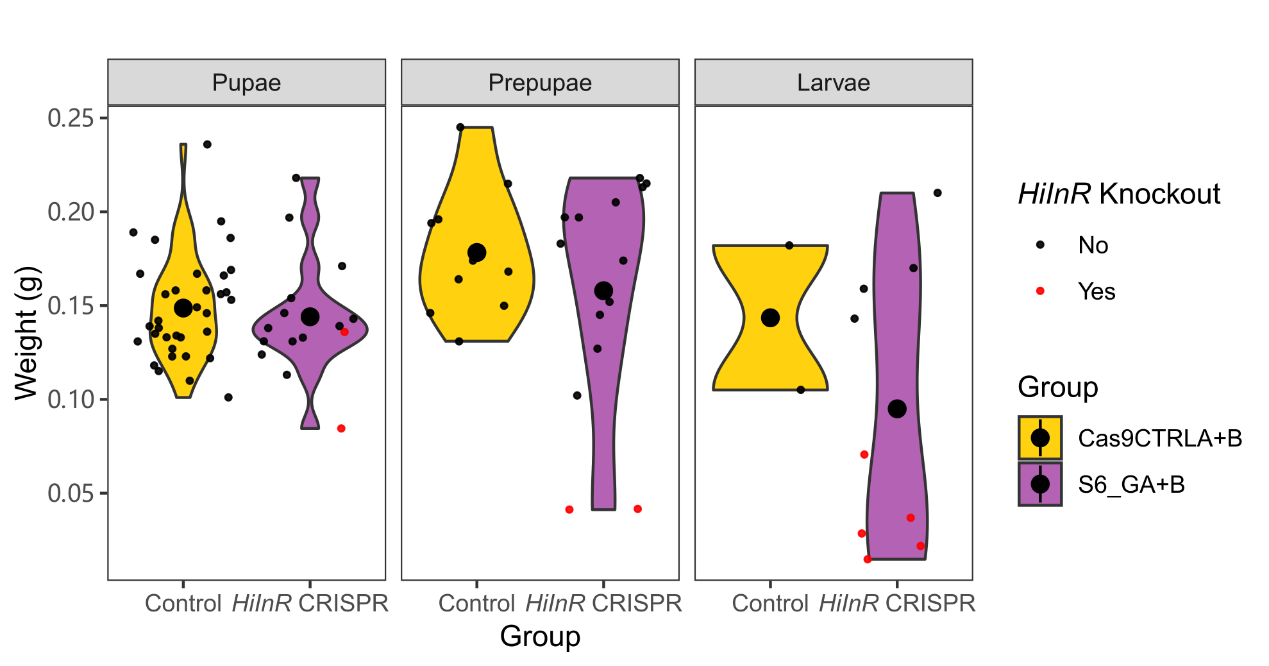

**Supplementary Figure 12** Violin plot for *HiInR* CRISPR individuals (purple, n=39) and all control individuals (yellow, n=47) at day 28 split by life stage (earliest being larvae and latest being pupae).

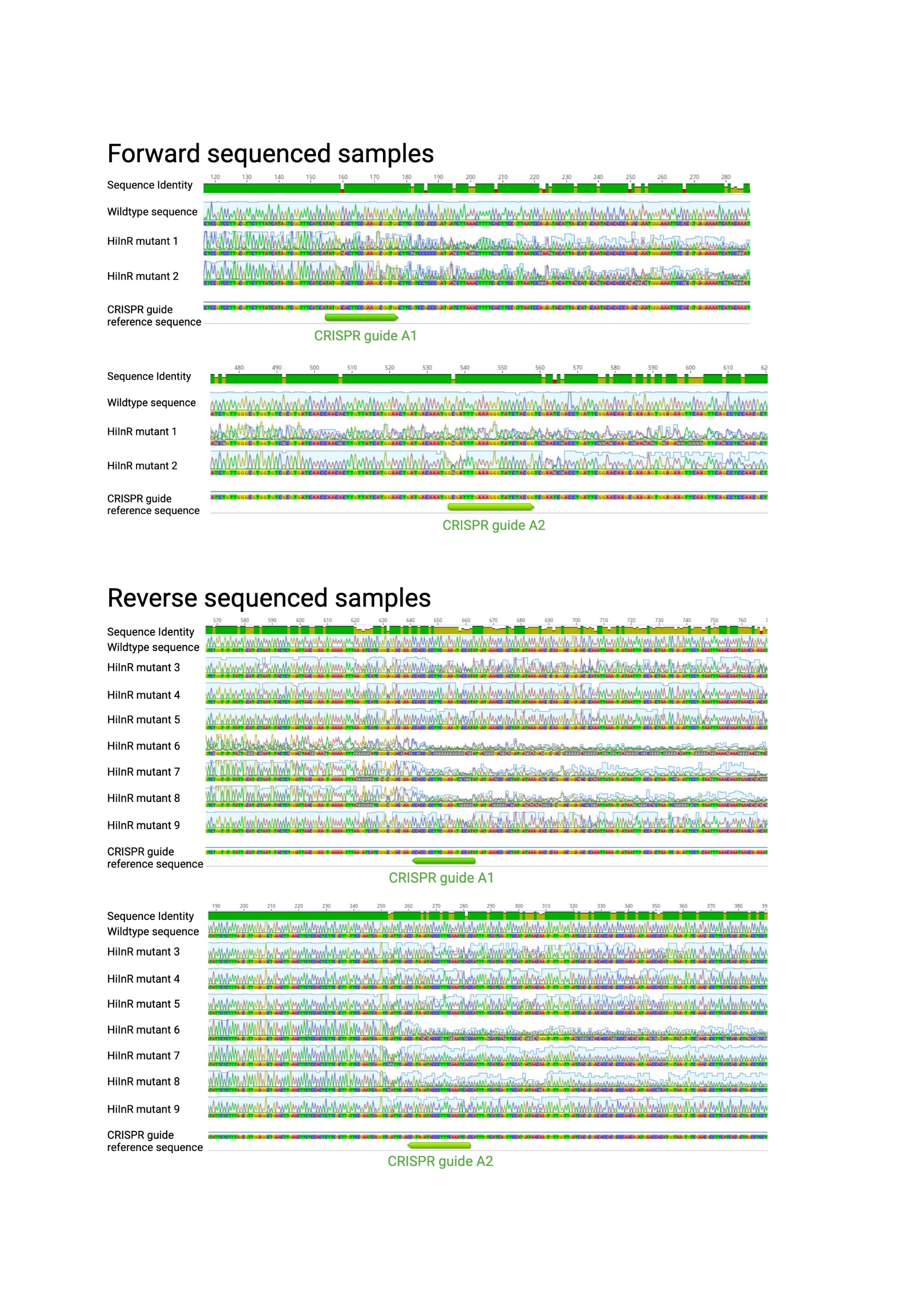

**Supplementary Figure 13** Sanger sequencing results of all mutants used in the study.
